# Real-time volumetric imaging of cells and molecules in deep tissues with Takoyaki ultrasound

**DOI:** 10.1101/2024.11.14.623368

**Authors:** Sunho Lee, Di Wu, Dina Malounda, Claire Rabut, Mikhail G. Shapiro

## Abstract

Acoustic contrast agents and reporter genes play a critical role in allowing ultrasound to visualize blood flow, map molecules and track cellular function in opaque living organisms. However, existing ultrasound methods to image acoustic contrast agents predominantly focus on 2D planar imaging, while the biological phenomena of interest unfurl in three dimensions. Here, we introduce a method for efficient, dynamic imaging of contrast agents and reporter genes in 3D using multiplexed matrix array transducers. Our “Takoyaki” pulse sequence uses the simultaneous scanning of multiple focal points to excite contrast agents with sufficient acoustic pressure for nonlinear imaging while efficiently covering 3D space. Through *in vitro* experiments, we first show that the Takoyaki sequence produces highly sensitive volume images of gas vesicle contrast agents and compare its performance with alternative imaging schemes. We then establish its utility in cellular imaging *in vivo* by visualizing acoustic reporter gene-expressing tumors in a mouse model of glioblastoma. Finally, we demonstrate real-time volumetric imaging by tracking the dynamics of fluid motion in brain ventricles after intraventricular contrast injection. Takoyaki imaging enables a more comprehensive understanding of biological processes by providing spatiotemporal information in 3D within the constraints of accessible multiplexed matrix array systems.

## INTRODUCTION

Ultrasound is a widely used technology for noninvasive biomedical imaging, valued for its deep penetration, high resolution, portability and safety. Recently, the traditional capabilities of ultrasound have been extended to large-scale visualization of biological systems and monitoring of cellular and molecular processes via volumetric 3D imaging^1^ and advanced contrast agents^2^. In addition to established synthetic microbubbles for vascular imaging^3^, emerging contrast agents such as nanobubbles^4^, phase-change nanodroplets^5^ and gas vesicles^6,7^ (GVs) make it possible to connect ultrasound with a larger variety of biological phenomena. GVs in particular have expanded the connection between ultrasound and biological function by serving as acoustic reporter genes^8–13^, molecular biosensors of enzymes and calcium^14,15^, and nanoscale contrast agents with enhanced circulation, extravasation, and biodegradation^16–21^.

In parallel, planar transducer arrays have made it possible to acquire 3D volumes at a single transducer position, facilitating dynamic and longitudinal imaging^22–29^. With decreasing costs due to advances in manufacturing and multiplexing, volumetric imaging is becoming increasingly practical and accessible.

Combining volumetric imaging with advanced contrast agents would greatly expand the capabilities of ultrasound in the biological and clinical settings by making it possible to visualize cellular and molecular phenomena in opaque volumes in real time. However, doing so carries significant challenges. Selective imaging of contrast agents relies on nonlinear pulse sequences that enrich contrast agent signals relative to background^30–33^. For example, GV imaging is typically performed with amplitude modulation (AM) methods^30,31^, which take advantage of reversible mechanical buckling of the GV under acoustic pressure to elicit nonlinear sound scattering distinct from mostly linear background^30,34,35^. Alternatively, BURST imaging detects GV expression with single-cell sensitivity by capturing distinctive signals produced transiently upon irreversible GV collapse^33^. Translating these approaches into 3D requires careful pulse sequence design within the constraints of array hardware.

Two classes of 3D imaging transducers compatible with financially accessible ultrasound systems (which typically have ≤ 256 electronic channels) are row-column arrays (RCAs) and multiplexed matrix arrays (MMAs). RCAs comprise two orthogonal, overlapping linear arrays with long elements^36–38^ and are capable of transmitting planar^27^ or sheet-like excitations^39^. Meanwhile, MMAs comprise a fully-addressed matrix probe (e.g. with 1024 elements) operated using a lower channel-count imaging system (e.g. with 256 channels) via an appropriate multiplexer (e.g. 4-to-1)^28,29,40^. While in principle MMAs provide greater flexibility than RCAs in their transmit and receive beam profiles, they carry their own constraints due to inter-element coupling and anisotropic element spacing, requiring creative pulse sequence schemes.

Here, we present a method for using matrix arrays to accomplish real-time volumetric imaging of acoustic contrast agents and reporter genes. Our “Takoyaki” paradigm uses all the elements of a square MMA to transmit focused ultrasound at multiple locations simultaneously using transmit delay functions resembling the Japanese street food Takoyaki (**Fig. 1a**). This approach shares the same philosophy as traditional multi-line transmission^41^, where a single transducer transmits multiple focused ultrasound beams at different steering angles simultaneously. Implemented on a 1024-element (32 × 32) MMA with a center frequency of 15 MHz and a 256-channel ultrasound acquisition system (**Fig. 1b-c**), Takoyaki imaging operates within the constraints of parallel element operation, while efficiently scanning the entire underlying volume with acoustic pressure suitable for nonlinear contrast imaging (**Fig. 1d**). In this study, we implement and optimize Takoyaki imaging and reconstruction, characterize its performance *in vitro*, and demonstrate its superiority relative to alternative pulse sequences compatible with MMA. We then illustrate its applications *in vivo* by selectively imaging brain tumors expressing acoustic reporter genes and monitoring fluid transport in brain ventricles in mice in real time using injected GV contrast. Our results establish Takoyaki imaging as a high-performance method for volumetric imaging of nonlinear contrast agents that is compatible with affordable ultrasound instrumentation.

**Figure 1:**
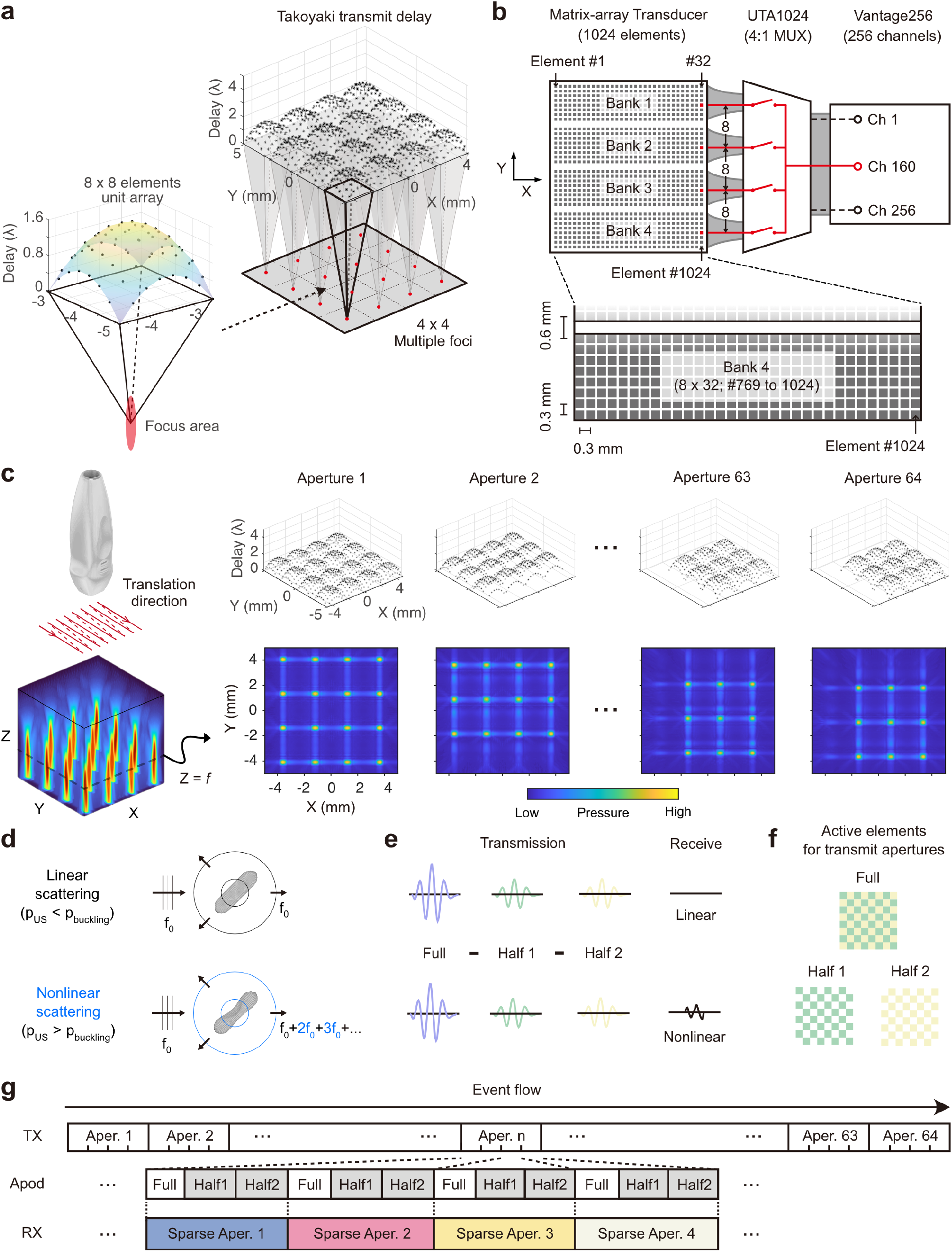
Takoyaki AM pulse sequence enables efficient scanning of 3D volume. **a**. Each unit array of 8 × 8 transducer elements generates focused ultrasound with a delay function concave in both X and Y, forming a cycle of Takoyaki functions. **b**. Equivalent diagram of the multiplexing system. The 1024 matrix-array transducer elements are sectioned by three inactive rows (gaps) into four banks (8 × 32 elements each). A 4-to-1 multiplexer connects one system channel with four elements of the matrix-array probe. These elements (red) operate in parallel within each bank, spaced by eight active elements. A single system channel can signal any subset of the four parallel elements, but delays and apodizations must be equal. **c**. Geometry of an element bank. **d**. The Takoyaki delay and its focused beams move along the X and Y axes. Top and bottom rows show the delay functions and simulated pressure fields at Z = focus (7 mm), respectively. **e**. Nonlinear response of GVs to the pressure above the buckling threshold. [adapted from ref.^31^] **f**. In AM, the echoes of two half-amplitude transmissions are subtracted from that of full amplitude to distinguish between linear and nonlinear scatterers. **g**. Two complementary checkerboard masks are used to generate half amplitude transmissions. **h**. Event flow of the Takoyaki AM sequence.

## RESULTS

### The Takoyaki sequence allows efficient acquisition of 3D volumes

In the Takoyaki sequence, an MMA transmits focused ultrasound at multiple locations simultaneously using Takoyaki-shaped delay laws, in which a hyperboloid function is periodically repeated across the 32 × 32 matrix array (**Fig. 1a, Fig. S1a**). Each hyperboloid, corresponding to a single focus, involves an 8 × 8 array of transducer elements. The 8-element period satisfies the parallel element operation constraints imposed by the 4-to-1 multiplexing system connecting the MMA to the ultrasound acquisition platform (**Fig. 1b-c**). To scan the entire volume, the focused beams are translated along both the X and Y axes, reconstructing the regions corresponding to the foci to cumulatively cover the underlying 3D field of view (FOV), with 64 different transmit apertures (**Fig. 1d**).

In the AM paradigm, the nonlinear response of GVs to ultrasound (**Fig. 1e**) is isolated from linear background scattering by subtracting the echoes of two half-amplitude transmissions from those of a full amplitude transmission (**Fig. 1f**). For Takoyaki AM imaging, the half-amplitudes are generated by applying two complementary checkerboard masks to the transmit apertures (**Fig. 1g**). The ultrasound imaging sequence involves transmitting 12 pulses (resulting from 4 receive apertures across 3 AM modes) for each transmit aperture. Signals are then received by a complementary set of four sparse random apertures^40^, and these are coherently compounded across all 1024 elements to generate a single volume (**Fig. 1h**). With a pulse repetition frequency (PRF) of 4 kHz, the acquisition of Raw-Frequency (RF) data for each complete volume image is achieved in 192 msec, calculated from the 250 µsec interval between transmissions multiplied by 64 transmit apertures with 12 transmissions each. To implement Takoyaki BURST imaging, the pressure is increased to collapse GVs, and GV signals are extracted by processing a temporal sequence of post-collapse frames.

Certain hardware considerations had to be addressed to put this sequence into practice. We implemented Takoyaki contrast imaging using a commercially-available 1024-element (32 × 32) MMA with a center frequency of 15 MHz (wavelength ∼ 100 µm) and an element pitch of 300 µm (lateral dimensions ∼ 1 cm × 1 cm). Being three times larger than the wavelength, this pitch is expected to generate substantial grating lobes, degrading image quality. However, with Takoyaki contrast imaging, we expect the pressure at the grating lobes to fall below the GVs’ nonlinear signal threshold, thereby suppressing grating lobe artifacts. A key benefit of the larger pitch is that it provides coverage of a 9-times larger FOV compared to wavelength-matched pitch for the same number of elements. Additionally, as with most MMAs, element placement is not isotropic, and there are gaps between element banks in the Y dimension (**Fig. 1b**). We took these gaps into account in delay calculation for each transmit aperture.

### Takoyaki imaging provides high quality isotropic contrast agent images

To examine the characteristics of the Takoyaki sequence, we imaged tissue-mimicking phantoms containing GVs using Takoyaki AM and compared these images with those obtained using two other potential imaging schemes compatible with MMA: sheet-pAM and 3D xAM (**Fig. 2a-b**). Sheet-pAM generates a sheet-like focus parallel to the XZ plane by cylindrically focusing ultrasound (**Fig. S1b, Methods**). In contrast, 3D xAM, a dimensional extension of xAM^31^ (recently implemented on RCA^39^), emits two counter-propagating axisymmetric angled plane waves and images their bisector slices parallel to the YZ plane (**Fig. S1c, Methods**). Both sequences form volume images slice-by-slice; on the contrary, the Takoyaki sequence forms volumes by reconstructing multiple lines in parallel. We did not consider Sheet-pAM and 3D xAM with alternate orientations (e.g. Sheet-pAM imaging YZ planes) since the gaps on the matrix array increase heterogeneity in Sheet-pAM’s pressure fields and disrupt the axisymmetry of the cross-propagating waves in 3D xAM. A sequence based on coherent plane-wave compounding was also not tested because its pressure levels are insufficient to robustly induce GV collapse for BURST imaging and may even be too low to drive AM-based imaging of certain GV types^10^ (**Fig. S2**). Prior to the experiments, we empirically optimized the transmission amplitudes for each sequence to maximize AM signals without GV collapse.

**Figure 2:**
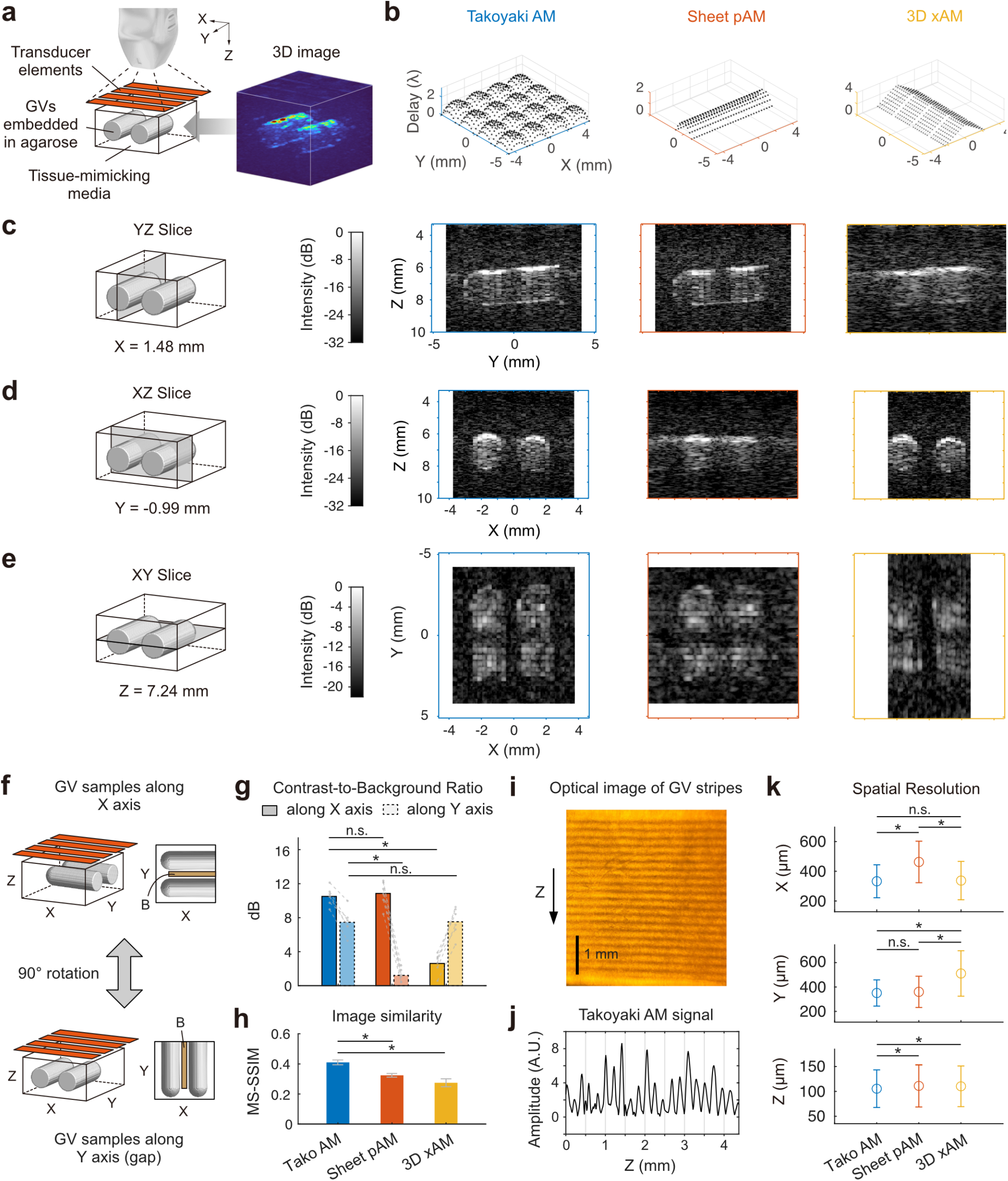
Takoyaki AM provides high quality isotropic images of gas vesicles. **a**. Experiment setup for imaging tissue-mimicking GV phantoms. The orange parallelograms represent the four banks of transducer elements. Each well in the phantom comprises a 5 mm length cylinder and a 1 mm radius hemisphere at one end, with an interspace distance of 1 mm. The axis of the wells was aligned with the Y axis, and the phantom location was adjusted so that its center was situated at Z = 7 mm. The 3D image of the phantom (OD = 6.5) acquired with Takoyaki AM is shown on the right. **b**. Representative delay laws of each sequence. **c-e**. Sliced phantom images acquired using different ultrasound sequences. The outer colored lines surrounding the sliced images indicate the maximum FOV. The color scale of the images is adjusted to match background levels across sequences. **f**. Imaging GV phantoms aligned along the X and Y axes. The brown box (B) represents the interspace background. **g**. CBRs of the volume images (N = 6). Solid and dashed lines correspond to phantoms aligned along the X and Y axes, respectively. **h**. Similarity analysis. **i**. Optical image of phantom with GV stripes. **j**. Takoyaki AM signal of GV stripes. **k**. Spatial resolution based on the full width half maximum. The error bars represent ±std (**h** and **k**). Asterisks denote p < 0.01. Two-sided paired t-test for **g** and **h**. Two-sided Mann-Whitney test for **k**. Takoyaki AM and Sheet-pAM transmissions were both focused at a depth of 7 mm.

Takoyaki AM demonstrated the highest isotropic image quality of the three sequences (**Fig. 2c-h**). Sheet-pAM and 3D xAM images were relatively less sharp and more susceptible to horizontal smearing artifacts along one axis (X and Y, respectively), likely due to their slice-by-slice beamforming (**Fig. 2c-e, Video 1**). To evaluate the impact of probe direction on each sequence’s image quality, we imaged the same phantom aligned along the X axis or rotated to lie along Y (**Fig. 2f**). When the phantoms were aligned along the Y axis, all three sequences showed some signal dropout around Y = 0 (**Fig. 2c,e, Fig. S3**), which we attribute to gaps in the matrix array and misalignment between transducer element banks^42^ (**Fig. S4**). Comparing their contrast-to-background ratios (CBRs) with the background measured in the interspace between the two GV wells, the CBRs for Sheet-pAM and 3D xAM were each significantly compromised along one direction (**Fig. 2g**). In comparison, Takoyaki AM maintained high CBRs in both orientations. Takoyaki AM images also exhibited the highest similarity after phantom rotation, as indicated by the multi-scale structural similarity (MS-SSIM) index^43^, further supporting its superior isotropic imaging capabilities (**Fig. 2h**).

To assess the spatial resolution of Takoyaki imaging and the two alternative sequences, we imaged phantoms containing GVs in stripe patterns of sub-wavelength thickness generated by acoustic patterning^44^ (**Fig. 2i-j, Fig. S5**). The measurements shared a similar tendency with the CBR analysis (**Fig. 2k, Table 1**): Takoyaki AM displayed the best spatial resolution for both lateral directions, while Sheet-pAM and 3D xAM showed poorer spatial resolutions in the X and Y axes, respectively. The axial resolution of Takoyaki AM was also better than those of Sheet-pAM and 3D xAM. With ∼330 μm lateral and ∼105 μm axial resolution, Takoyaki imaging meets expectations based on element pitch and transmit wavelength.

**Table 1:** Spatial resolution of Takoyaki AM, Sheet-pAM, and 3D xAM.

| Unit: $\mu\text{m}$<br>(mean ± std) | X | Y | Z |
| --- | --- | --- | --- |
| Takoyaki AM | $333 \pm 111$ | $351 \pm 107$ | $105 \pm 37.7$ |
| Sheet-pAM | $463 \pm 140$ | $360 \pm 128$ | $111 \pm 42.3$ |
| 3D xAM | $337 \pm 129$ | $511 \pm 186$ | $110 \pm 40.8$ |

All three imaging modes cover a partial FOV underneath the array (**Fig. 2c-e**). Takoyaki AM provides a FOV area (7.5 × 8.4 = 63 mm^2^) between those of 3D xAM (5.1 × 10.2 = 52.02 mm^2^) and Sheet-pAM (9.3 × 8.4 = 78.12 mm^2^).

As a result of its combined high isotropy, CBR, resolution and FOV, Takoyaki AM allows the matrix array probe to produce high-quality volumetric images of contrast agents. In addition to these performance advantages relative to alternative pulse sequences, Takoyaki AM’s two-dimensional parabolic focusing can, in principle, achieve higher pressures compared to 3D xAM and plane-wave-based sequences, making it more suitable for imaging modes that require exceeding specific pressure thresholds, such as BURST imaging.

### Takoyaki imaging enables volumetric visualization of genetically labeled brain tumors

To examine if the Takoyaki ultrasound sequence can selectively image genetically labeled cells in 3D *in vivo*, we tested it using a brain tumor model. The ability to image brain tumors *in situ* is critical to understanding their growth dynamics, interactions with the neurovasculature and healthy brain tissue, and response to therapeutic interventions. Doing so in 3D would provide a more complete picture of these interactions and facilitate acquisitions that are less operator-dependent. We engineered U87 human tumor cells to express GVs as acoustic reporter genes and implanted them into mouse brains^45^ (**Fig. 3a**). These tumor cells were designed to form GVs and produce ultrasound contrast upon induction with the small molecule doxycycline. Following 6 days of tumor progression and a 2-day induction period, the skull was removed to enable acquisition of Takoyaki AM images of the brain (**Fig. 3b**). During this imaging session, power Doppler images were also acquired with the MMA system^29^ to provide vascular context to the tumor images, along with Takoyaki B-mode images for enhanced anatomical reference. For comparison, a 3D reconstruction of the tumors was generated by systematically translating a 15 MHz linear array probe (0.1 mm pitch) in 0.2 mm intervals and artificially stacking the resulting 2D xAM^31^ images (denoted as st-xAM image).

**Figure 3:**
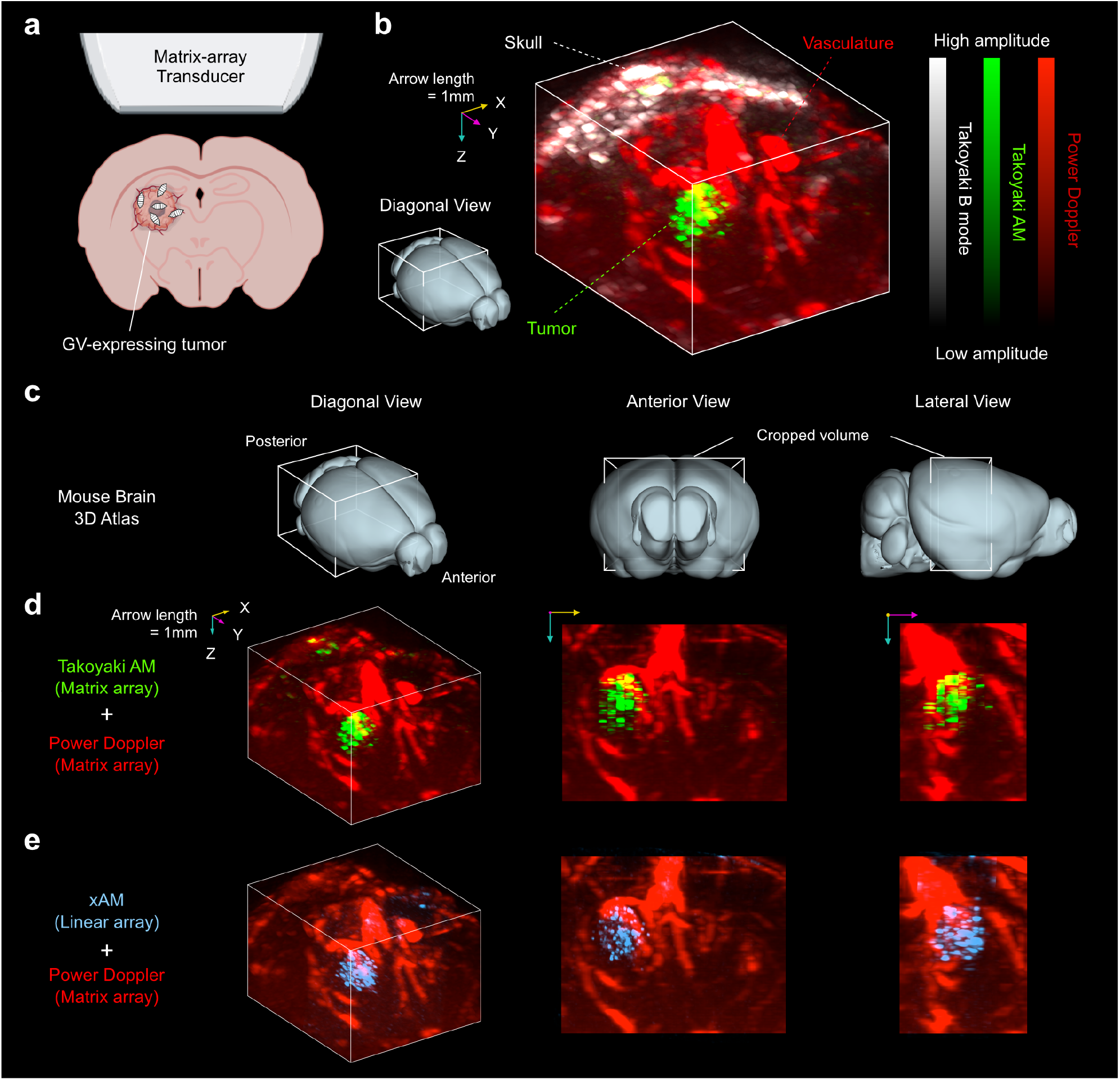
Takoyaki AM enables volumetric imaging of genetically labeled tumors in the mouse brain. **a**. After tumor cell injection and growth, we induced GV expression and performed ultrasound imaging. Created in BioRender. **b**. Overlay of Takoyaki B-mode (white), Takoyaki AM (green), and power Doppler images (red), acquired using an MMA transducer. **c**. The Allen Mouse Brain Atlas^46^ in the top row illustrates brain orientation, with the white box indicating the imaging regions. **d**. Takoyaki AM **e**. Stacked xAM images obtained with a sweeping linear array probe (blue). For the anterior and lateral views, a half-cropped 3D image is used to exclude the skull side. Colored arrows represent the axes, with lengths of 1 mm.

Takoyaki AM successfully provided selective imaging of brain tumors, capturing their 3D volume and location inside the brain (N = 2; **Fig. 3c-d, Fig. S6**). Aligning Takoyaki AM with power Doppler images enabled precise localization within the vascular anatomy (**Fig. 3d, Video 2**). While st-xAM achieved similar results (**Fig. 3e, Video 2**), it required mechanical scanning. CBRs were 15.27 dB for Takoyaki AM and 15.22 dB for st-xAM, indicating comparable clarity. The spatial consistency between the two modes was high, though some minor regions detected in st-xAM were missed by Takoyaki AM, likely due to array signal gaps. Overall, these findings confirm Takoyaki imaging’s capability to visualize brain tumors through GVs *in vivo*.

### Takoyaki AM enables dynamic real-time imaging of nanoparticle transport in brain ventricles

The matrix probe’s ability to perform rapid volumetric scanning makes it suitable for observing dynamic processes within a volume. Leveraging this feature, we explored the feasibility of real-time monitoring of GV dynamics within the brain ventricles. The ventricles are filled with cerebrospinal fluid (CSF), whose circulation plays a vital role in clearing metabolic waste and transferring nutrients and drugs to brain tissue, especially during sleep^47–49^. Being able to visualize the flow of CSF and particles contained within it is critical for comprehending its normal biological function and involvement in various diseases^48,49^. In addition, understanding CSF transport can inform work on drug delivery to the brain, in which therapeutics such as viral vectors are administered into the ventricles to circumvent the blood-brain barrier^50,51^.

To demonstrate dynamic imaging of induced CSF flow, we injected GVs into one of the brain ventricles and followed their distribution in real time (**Fig. 4a**). While real-time imaging, we infused 4.5 *μ*L of GV solution (1.6 nM) into the right lateral ventricle (LV) at a rate of 75 nL/s over 60 sec. Before, during and after the infusion, we continuously acquired volume images, choosing a frame rate of 0.26 Hz (1 frame per 3.8 sec) to capture the expected dynamics and allow real-time image processing and display. This frame rate can be accelerated to >5 Hz by deferring or accelerating image reconstruction and storage (currently ∼2.8 sec/frame and ∼1 sec/frame, respectively). Power Doppler images were acquired before the GV injection as an anatomical reference. To quantify the dynamics of GV transport across the ventricles, we outlined them (**Fig. 4b, Fig. S7a**) and measured their AM signal density over time (**Fig. 4c, Fig. S7b**).

**Figure 4:**
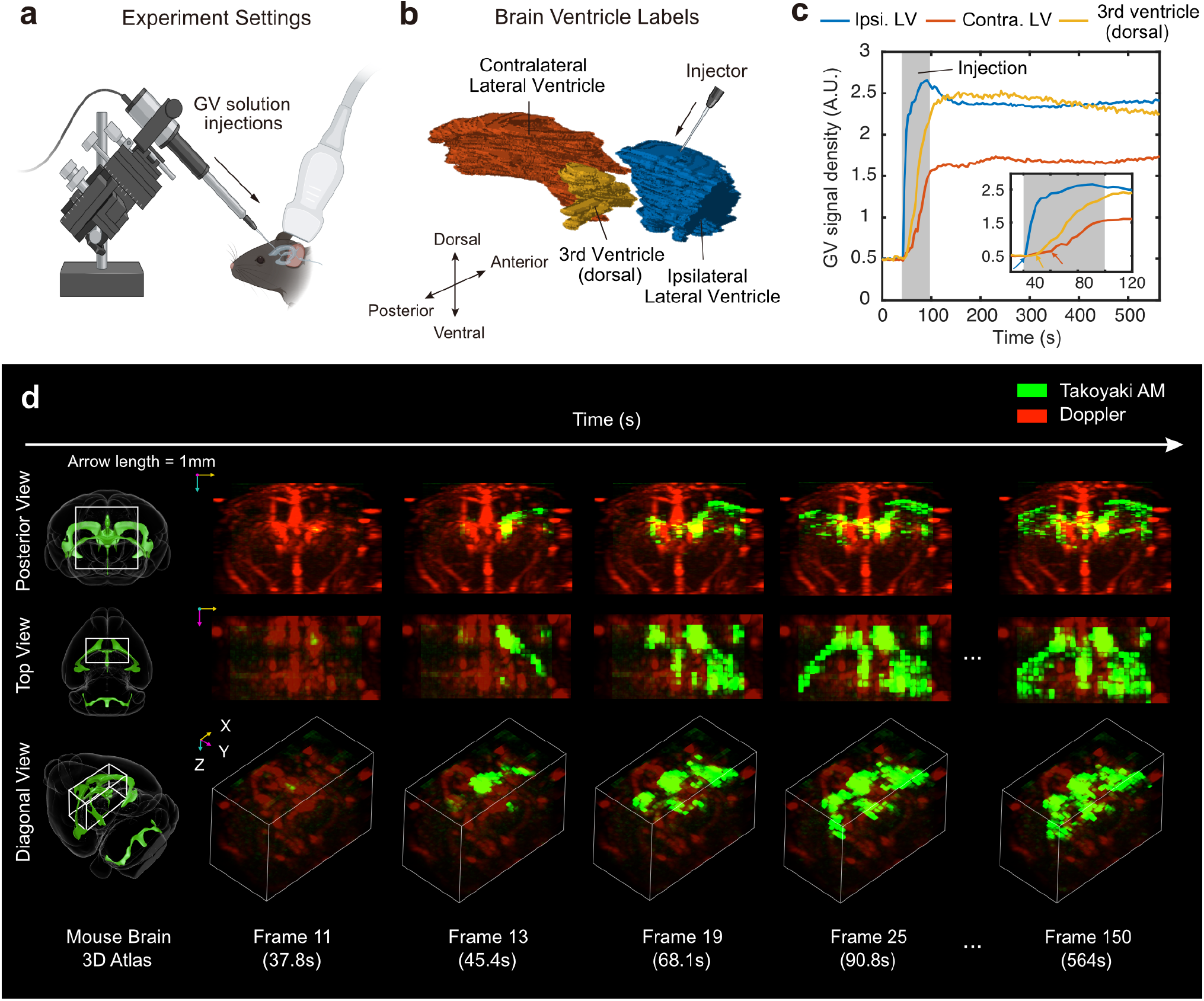
Takoyaki AM enables real-time dynamic imaging of nanoparticle transport within mouse brain ventricles. **a**. Microinjection of GV solutions into the right LV. Created in BioRender. **b**. Ventricle boundaries outlined based on GV signals. Blue, red, and yellow denote the ipsilateral LV to the injection site, contralateral LV, and third ventricle, respectively. Outlines were drawn manually based on images. **c**. Temporal evolution of GV signal density. Injection commenced between Frame 11 and 12 (∼40 sec), and lasted for 60 sec, as indicated by the gray shaded area. The arrows in the inset mark the start of the rapid signal increases for each ventricle. **d**. Selected frames from stored real-time images. The Allen Mouse Brain Atlas^46^ in the leftmost column illustrates brain orientations, with white bounding boxes indicating the image location. Colored arrows have a length of 1 mm along their corresponding directions. The sampling interval was approximately 3.8 sec.

During real-time monitoring, we successfully captured the ventricular system and the propagation of GVs across both LVs and the third ventricle (N = 2; **Fig. 4d, Fig. S7c, Video 3**), clearly distinguishable from the vascular structures seen in Doppler images. Initial GV dispersion was observed in the LV ipsilateral to the injection site, followed by distribution to the third ventricle and then to the contralateral LV. The signal density graphs reflected the nanoparticle spreading dynamics observed during the real-time recording (**Fig. 4c, Fig. S7b**). Rapid signal increases, corresponding to GV influx and spreading, appeared sequentially in the ipsilateral LV, third ventricle, and contralateral LV. These results demonstrate the ability of Takoyaki AM imaging to track a dynamic biological process in 3D, representing the first time to our knowledge that cerebrospinal fluid transport has been visualized with ultrasound.

### Takoyaki BURST enables higher-sensitivity volumetric imaging in gap areas and deeper regions

In addition to its capacity for AM imaging, the Takoyaki sequence is well suited for ultrasensitive imaging using BURST^33^, which requires higher transmit pressure to irreversibly collapse GVs and isolate the resulting strong, transient signal. To validate its efficacy, we used the Takoyaki BURST sequence (**Fig. 5a**) to acquire images of brain tumors and ventricles following the completion of the corresponding AM imaging sessions.

**Figure 5:**
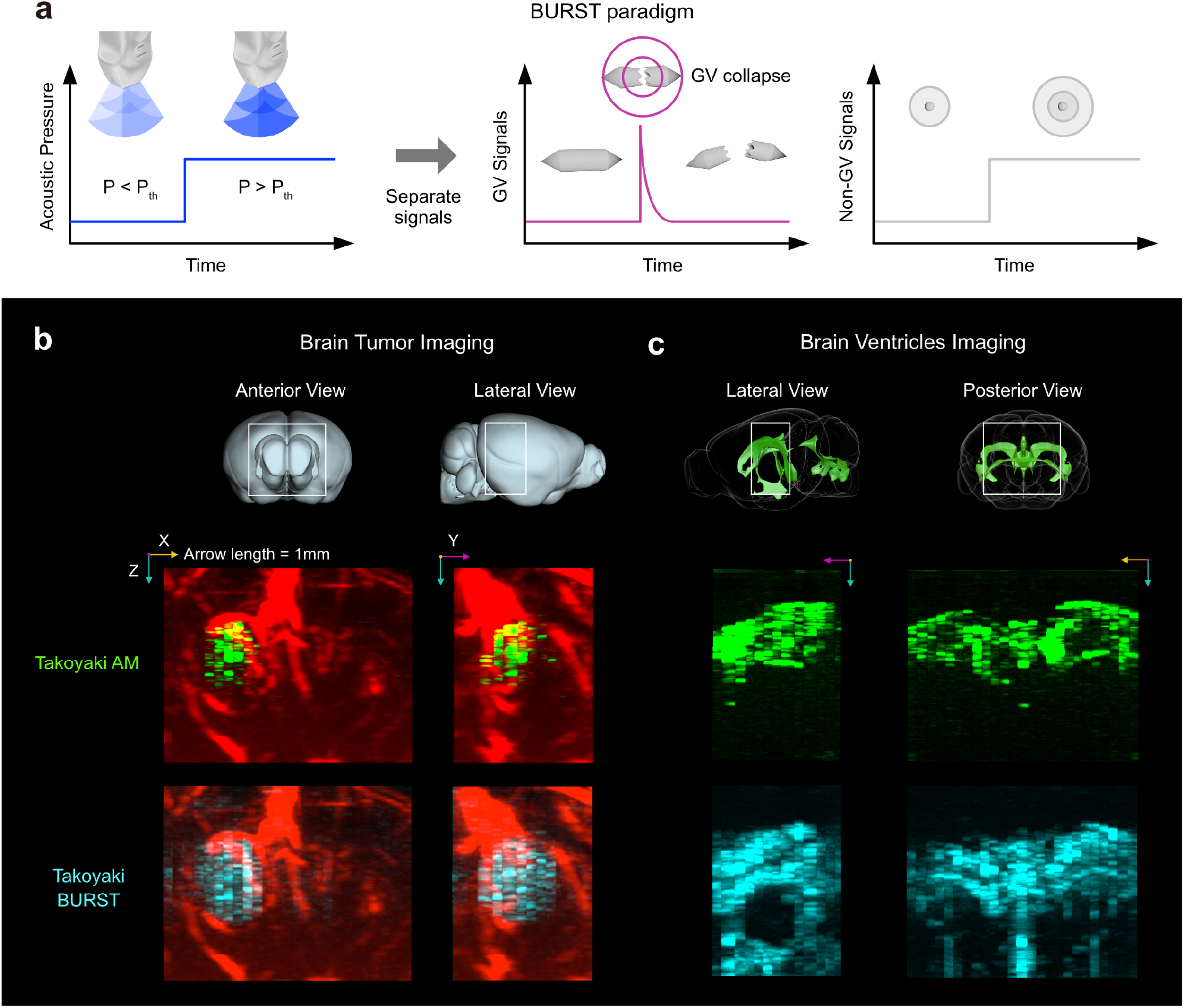
Takoyaki BURST enables higher-sensitivity volumetric imaging in gap areas and deeper regions. **a**. BURST paradigm. Application of pressure exceeding the collapse threshold (P_th_) induces GV collapse, leading to strong transient GV signals, while non-GV signals remain constant. The two signals are separated after acquisition through processing. **b, c**. Comparison between Takoyaki AM (green) and BURST (cyan). The Allen Brain Atlas^46^ in the top row illustrates brain orientations, with white bounding boxes indicating the imaging regions. Colored arrows representing axes are scaled to a length of 1 mm along their respective directions. **b**. Anterior and lateral views of a brain tumor. Doppler images (red) are overlaid. **c**. Lateral and posterior views of brain ventricles.

In brain tumor imaging, our Takoyaki BURST sequence demonstrated enhanced visualization capabilities, effectively capturing signals from gap areas in the array and deeper regions that were difficult to discern with AM alone (**Fig. 5b, Fig. S8a, Video 4**). This improved depth sensitivity was especially beneficial for imaging brain ventricles, where

BURST clearly visualized the ventral portion of the third ventricle and LVs (**Fig. 5c, Fig. S8b, Video 4**), whereas AM imaging was limited to dorsal regions. While optimization of GV expression levels, injected concentration, and focal beam parameters could enhance Takoyaki AM performance, these results show that the 3D BURST approach offers a robust way to maximize signal detection across the volume when a single snapshot is sufficient.

## DISCUSSION

In this work, we introduced Takoyaki volumetric ultrasound sequence, an imaging approach that enables efficient capture of contrast agent and reporter gene signals in 3D using the emerging class of commercially available MMA transducers. Using a repeated hyperbolic delay pattern that adheres to the parallel element constraints of an MMA, the Takoyaki sequence maximally utilizes the available transducer elements to generate focused transmissions at multiple locations simultaneously. This provides efficient scanning of the underlying volume while applying sufficient transmit pressure to elicit robust nonlinear contrast signals. Using this approach to implement nonlinear AM and BURST imaging, we showed that Takoyaki imaging can effectively capture the 3D distribution of specific cells (brain tumors) and biomolecules (gas vesicles) inside opaque tissues such as the brain. Furthermore, Takoyaki AM enables dynamic 3D imaging, revealing for the first time the real-time transport of nanoparticles across brain ventricles. Meanwhile, Takoyaki BURST provided highly sensitive and comprehensive 3D snapshots of contrast agent distribution. Compared to alternative pulse sequences we implemented on MMA, Takoyaki imaging provided the best combination of sensitivity and isotropy.

Given the relative accessibility of 256-channel ultrasound scanners and MMAs, and the global availability of GV-based genetic constructs^10,52^, we anticipate the use of Takoyaki imaging in a wide range of biological research and biomedical applications. In addition, Takoyaki AM should be applicable to any other ultrasound contrast agent with nonlinear pressure responses, such as microbubbles^2^ and nanobubbles^4^, and extensible to other imaging modes such as pulse inversion. Where higher pressures are needed, they could be amplified by distributing power across fewer banks of the array during transmission. For instance, a (4 × 1)-form Takoyaki BURST, using only one out of four parallel elements, could generate approximately double the pressure of the original (4 × 4)-form.

The presented Takoyaki approach can be improved with future modifications. The heterogeneity of pressure fields caused by transducer element misalignment and gaps leads to loss of signal in certain regions of the FOV. This pressure heterogeneity could be mitigated by adjusting the apodization amplitude for each transmission. Furthermore, imaging depth could be improved by implementing multi-depth delays and managing contrast agent concentrations to avoid shadowing. Dynamic imaging with Takoyaki AM can already be fast enough for many biological phenomena (192 msec per frame), but is currently slowed down by data processing and storage steps when real-time display is required. Online reconstruction could be accelerated by implementing it on graphical processor units. In theory, Takoyaki AM could ultimately reach an acquisition speed of 17.4 msec per frame for a 1-cm imaging depth, considering the minimum pulse-echo time of flight. With these enhancements and options, Takoyaki imaging will come in multiple flavors to satiate any hunger for 3D contrast-enhanced, biomolecular and cellular ultrasound.

## MATERIALS AND METHODS

### Ultrasound acquisition system

The 15 MHz matrix array probe (Vermon) comprises 35 × 32 transducer elements with a 0.3 mm pitch size. The elements are organized into four active banks, each containing 8 × 32 elements, separated by three inactive rows (**Fig. 1b-c**). The matrix probe was connected to a programmable ultrasound system (Verasonics Vantage 256) via a multiplex adapter (Verasonics UTA 1024-MUX). Each system channel can control up to four elements (i, i + 256, I + 512, i + 768) in parallel, where i ∈ {1, 2, …, 256}, mandating that elements at equivalent positions in each bank share the same delay and apodization settings. A waveform with a 15.625 MHz center frequency, 0.67 duty cycle, and 2 half-cycle periods was used, unless specified otherwise. All ultrasound sequence scripts were written and implemented via MATLAB 2022a (MathWorks).

### Takoyaki AM/B-mode sequences

The Takoyaki-like delays use unit arrays of (8 × 8) transducer elements to create focused ultrasound beams. The first transmission aperture, representing the basic form in the Takoyaki sequence, consists of 16 unit arrays arranged in a 4 × 4 grid, forming a full aperture of (32 × 32) elements (**Fig. 1a**). Throughout the transmission session, this basic delay pattern is translated across the entire transducer elements. There are 8 possible translations along both the X and Y axes – in total, 64 different apertures. This translation is akin to moving the matrix array probe across the XY plane while maintaining the basic Takoyaki delay pattern (**Fig. 1d**). Note that changes in gap locations within the unit arrays were considered in the delay calculations. The imaging regions for each transmission are basically cuboids (Frustum-shaped volumes of 0.3 mm × 0.3 mm x depth range) centered at each focus; however, for the 2^nd^ and 8^th^ translations along the Y direction, the Y width of the cuboids was set to 0.45 mm instead of 0.3 mm to compensate for the longer jump caused by gaps between transducer elements, resulting in the cuboids not being exactly centered at the foci. We inactivated elements on the unit arrays that are truncated after translating the delay function. Although this leads to a change in the number of parallel elements used and thus variations in pressure levels for different aperture types (8 apertures use all 4 parallel elements, and the others use 3 parallel elements), we found that for our matrix array probe, this inactivation gave less variance in pressure levels at the focal area across different aperture types compared to not performing such inactivation.

In the AM sequence, a full-amplitude ultrasound transmission is followed by two half-amplitude transmissions, and the received signals corresponding to the half-amplitudes are subtracted from that of the full-amplitude (**Fig. 1f**). Conversely, B-mode imaging uses only the full-amplitude transmission. For the Takoyaki AM sequence, the half-amplitude ultrasounds are generated by applying two complementary checkerboard masks to the transmission apodization (**Fig. 1g**). For the receive apertures, a complementary set of four sparse random apertures^40^ was used. Each transmission aperture undergoes an AM sequence (three transmissions) for each of the four receive apertures, and the received signals are coherently compounded (**Fig. 1h**). Data transfer and reconstruction occur once RF data for all 64 transmit apertures are collected. After the reconstruction, the image is visualized and then stored. The focal length was 7 mm. The reconstruction depth of interest was set from 3.3 mm to 10 mm, with a voxel size of half wavelength (λ/2 = 49.3 *μm*) for all sides. The pulse repetition frequency (PRF) was 4 kHz. Pressure field simulations were performed using the TXPD tool within the Verasonics Vantage software.

### Takoyaki BURST sequence

In the BURST paradigm^33^, a pressure higher than the collapse threshold of GVs is applied following a lower pressure, and distinctive transient GV signals are then extracted from the background (**Fig. 5a**). We used two different sequences to obtain the BURST images.

First, we developed an independent script that implements the BURST scheme without any coherent compounding. This script is fundamentally based on the event flow of Takoyaki B-mode but employs only one receive aperture per transmit aperture type. Additionally, the receive aperture for each transmit aperture varies randomly and is not identical. The number of frames captured at low pressure (Voltage = 1.6 V, Peak positive pressure = 60 kPa) was set to 2, while at high pressure (Voltage ≥ 27 V, Peak positive pressure ≥ 2 MPa), it was set to 28. Unlike the live imaging (Takoyaki AM) script, which reconstructs images after obtaining the RF data for each frame, this BURST script saves the RF signals for all frames first and then reconstructs the images later. IQ data are accumulated across frames. The PRF and time interval between frames were 4 kHz and 20 msec, respectively. The focal length was 7 mm. The number of half cycles was set to 3. This script was used for brain tumor experiments.

Alternatively, for brain ventricle imaging experiments, we utilized the live imaging script of the Takoyaki sequence with a high-pressure setting (Voltage ≥ 28 V). After increasing the pressure, ten frames were stored and then processed to extract the GV signals. The same parameters used in the Takoyaki AM imaging were applied.

### Sheet-pAM and 3D xAM

In Sheet-pAM, each transmission involves an array of (32 × 8) transducer elements, generating a sheet-like focus parallel to the XZ plane (**Fig. S1b**). During imaging, a (9.6 mm × 0.3 mm x depth range) cuboid region centered at the focus is reconstructed. The focus moves across the entire array along the Y axis through 25 sequential translations. To compensate for larger jumps due to gaps in the matrix array, the same method used in Takoyaki AM was applied. The focal length was 7 mm.

For 3D xAM, each transmission utilizes an array of (16 × 32) elements. Each half (8 × 32) emits an angled plane wave that is parallel to the Y axis and symmetric with respect to the other (**Fig. S1c**). The bisector slice (0.3 mm × 10.5 mm x depth range) parallel to the YZ plane is reconstructed, similar to xAM with a linear array^31^. The X-waves were angled at 6° relative to the surface of the matrix array probe and translated 17 times along the X axis.

Both AM sequences share the same event flow structure as Takoyaki AM, but 3D xAM differs in how the half-amplitude ultrasound is generated. In 3D xAM, the half-amplitude ultrasound is produced by using only each half (8 × 32) elements, whereas Sheet-pAM employs checkerboard masks on its apertures. The depth range and other parameters were identical to those used in Takoyaki AM. Pressure field simulations were performed using the TXPD tool within the Verasonics Vantage software.

### BURST processing algorithm

BURST signals are traditionally processed using the temporal template unmixing algorithm^33^, which assumes that the peaks of GV signals only appear in the frame immediately following an increase in pressure. For example, the template vector for GV signals is defined as *u*_*g*_ = [0 1 0 0 0 0]^*T*^, when the first frame corresponds to low pressure and the subsequent frames to high pressure. In contrast, non-GV signals remain constant after increasing pressure, which is represented by the template vector *u*_*s*_= [0 1 1 1 1 1]^*T*^. However, in our cases, we observed that GV collapse occurred over multiple frames, seemingly progressing towards deeper areas, possibly due to a shielding effect. Moreover, since we accumulated IQ data across frames, the background signal did not remain at the constant level but showed an increasing trend. Therefore, we used alternative methods for processing BURST signals instead of relying on the template unmixing algorithm.

Inspired by Demené *et al*.^53^, we took advantage of Singular Value Decomposition (SVD) to filter out coherent background signals from GV signals. Let *S* ∈ ℝ^*M×N*^ denote the spatiotemporal matrix formed by reshaping the N frames (4D matrix) into a 2D matrix. Then, SVD decomposes S as follows,

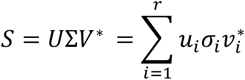

where *u*_*i*_is the *i*^th^ column of the matrix U, *v*_*i*_ is the *i*^th^ column of the matrix V, σ_*i*_ is the *i*^th^ singular value, and *r* is the rank of *S*. SVD can be effective in our context, where the exact trend of the background template vector is uncertain and there is minimal movement across frames. This is because the Eckart-Young theorem^54^ guarantees that the multiplication of the first components of SVD,

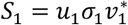

provides the best rank-1 approximation of the spatiotemporal matrix *S*. Given that the background signals can be interpreted as a spatial vector multiplied by a global temporal vector, subtracting *S*_1_ from *S*, denoted as *S*−*S*_2_, would effectively remove the background (linear scatterers), assuming that GV signals are sufficiently localized. Filtering out the first components of SVD can be expressed as,

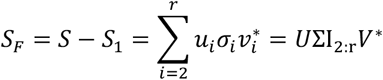

where I_2:r_ ∈ ℝ^r×r^ is a modified identity matrix with its first diagonal element set to zero. Although additional noise reduction may be achievable by discarding some of the last SVD components, we chose not to implement it for the sake of simplicity. Assuming successful background removal from the spatiotemporal images and no other sources of strong fluctuations,

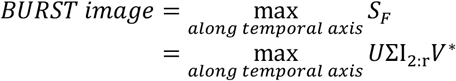

could properly extract peaks from GV collapse signals along the temporal axis. We excluded pre-collapse frames as their inclusion in *S* degraded the quality of BURST images. (In the case of the temporal unmixing algorithm, the insignificance of pre-collapse frames for calculating BURST signals can be mathematically proven, although we omit the proof here for brevity.)

### Doppler imaging sequence

We developed the Doppler imaging sequence by applying the 3D ultrafast Doppler imaging paradigm^23^ to the “Direct” multiplexing combination^29^ of transmit (TX) and receive (RX) banks. In this sequence, for each steering angle, one TX bank transmits a tilted plane wave, and the corresponding RX bank from the direct combinations receives the signals. This process is repeated for each of the four banks. We used six different steering angles: [0°, 0°], [2.5°, –2.5°], [-2.5°, 0°], [2.5°, 2.5°], [0°, –2.5°], and [0°, 2.5°]. Therefore, a total of 4 TX/RX pairs x 6 angles = 24 events were combined through coherently compounding to form a single volume. A power Doppler image was calculated using 200 volume acquisitions with a PRF of 26316 Hz (38 μsec between pulse transmissions).

Data transfer and beamforming followed the super-volume methods described in Yu *et al*.^28^. The RF data from the 200 volume acquisitions were packed into four blocks (or super-volumes), each stored in a different receive frame. Data transfer to the host occurred after each block was obtained. Once data acquisition and storage were complete, beamforming was performed using a separate script specialized to reconstruct IQ data. SVD^53^ was then applied to filter the spatiotemporal IQ data before processing them into a power Doppler image. The voxel size of the reconstructed images was equal to the wavelength for all sides. The FOV of the doppler images was 9.6 × 9.6 mm.

### GV preparations

As explained in Lakshmanan *et al*.^55^, after growing the Anabaena cells, GVs were harvested from floating Anabaena cells. GVs were released from the cells by hypertonic lysis and then purified by using repeated buoyancy-assisted centrifugation and resuspension. GvpC removal was performed by using GV stripping buffer, which contains urea, and by repeated buoyancy-assisted centrifugation and resuspension. The concentration of GVs was measured based on optical density (OD) at 500 nm using a spectrophotometer (Nanodrop 2000c).

### Contrast-to-background ratio (CBR) calculation for in vitro experiments

Phantoms consisted of cylindrical wells filled with GVs embedded in 0.5% agarose, surrounded by a background media made from 0.2% 3 μm Al^2^O^3^ + 1% Agarose. To prepare the GV mixture, GV solutions with an OD of 13 were mixed with 1% agarose in a 1:1 ratio, resulting in a final OD as 6.5. The phantoms (N = 6) were submerged in a water bath and imaged with their cylindrical wells aligned parallel to the X axis (**Fig. S3c-e**) and rotated to be parallel to the Y axis (**Fig. 2c-e**). XY slices were visualized in **Fig. 2e** and **Fig. S3e** by averaging five adjacent slices. The order of the three ultrasound sequences and the two orientations was permuted for each phantom. We empirically optimized the transmission amplitudes for each sequence to maximize AM signals without GV collapse.

The CBR of the GV phantoms was calculated using the following equation,

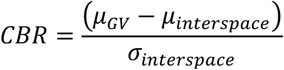

where *μ*_*GV*_ is the average signal intensity of the two GV wells, *μ*_*interspace*_ is the average signal intensity of the center interspace between the wells, and σ_*interspace*_ is the standard deviation of signal intensities in the interspace. A two-sided paired samples t-test was conducted to compare the CBRs between each pair of different sequences.

To define the regions corresponding to the GVs and the interspace, we first created a volumetric mask based on the geometry of the phantoms, and manually registered it to each volumetric image. The mask was then eroded using a sphere with a radius of 5 voxels (2.5 times the wavelength).

The interspace region was defined as a rectangular box situated between the cylindrical regions of the two GV wells, positioned 7 voxels away from the unshrunk mask of each GV well (**Fig. 2f**). The cylindrical regions were defined where both wells of the eroded masks exhibited the maximum circular area in a slice. The range along the Z axis was determined by the overlap of two unshrunk wells along the Z axis, further reduced by 7 voxels from the upper end.

When the phantom was aligned parallel to the X axis, two additional steps were taken. First, because of the distortion of images along the Y axis, the distance between the two wells appeared shorter than the actual ground truth. To account for this, the volumetric mask was adjusted based on this shortened distance. For phantoms aligned along the Y axis, the length of the mask was similarly adjusted. Second, since the mask was truncated by the FOV of 3D xAM (the only case of truncation, **Fig. S3e**), we applied the ROIs from 3D xAM images to the corresponding images from the other sequences.

### Image similarity analysis

For each phantom, we obtained transformation matrices between the volumetric masks of the phantom aligned along the X axis (mask X) and the Y axis (mask Y). Specifically, using the Medical Registration Estimator application in MATLAB R2024b (MathWorks), we first manually registered mask Y to the fixed mask X, and then performed automatic deformable registration. After confirming that the registration was successful (structural similarity (SSIM) index ≥ 0.99), we stored the affine transformation matrix and the displacement field produced by these steps.

Next, we applied the transformation matrices to the images of the corresponding phantoms aligned along the Y axis and calculated the multi-scale SSIM (MS-SSIM)^43^ between the transformed images and the images of phantoms aligned along the X axis. The region of interest was set to the bounding box of mask X, and the range along the X axis was further reduced to match the FOV of 3D xAM. The intensity of all images was compressed using a base-10 logarithm before computing their MS-SSIM values. A two-sided paired samples t-test was conducted to compare the MS-SSIM values between each pair of different sequences.

### Spatial resolution measurements

To measure the spatial resolution of the ultrasound sequences, we used phantoms (N = 6 for both axial and lateral resolution measurements) with GVs embedded in 0.5% agarose gel, arranged in a stripe pattern with subwavelength-scale widths. Solidified GV mixtures with the desired OD (OD = 2 for axial resolution, and OD = 3 for lateral resolution) were first prepared by mixing GV solution with 1% agarose in a 1:1 ratio. Next, two different methods were employed to create the stripe patterns for axial and lateral resolution assessments.

For axial resolution (Z axis), phantoms with stripe intervals of 0.2 mm and widths of ∼100 μm were created using acoustic patterning^44^, following the method used in Rabut *et al*.^32^ (**Fig. S5a**). For lateral resolution, we utilized the elevational focusing of a linear array probe (Verasonics, L22-14vX) to generate a sawtooth pattern of GVs with a 1 mm interval. This was achieved by repeatedly moving and collapsing the GVs using plane wave transmissions (**Fig. S5b**). We then selected an XY slice as close to the top (thinnest part) of the sawtooth as possible, ensuring that there were still sufficient GV signals. The depth of the extracted slice was Z = ∼7 mm. The voxel sizes of the reconstructed images were set to (X, Y, Z) = (1, 1, 1/8) wavelength for axial resolution measurements and (1/2, 1/2, 1/2) wavelength for lateral resolution. Transmission amplitudes were empirically optimized for each sequence to maximize AM signals without GV collapse. The optical images of these phantoms were taken with a microscope, transmitting a bright light.

For peak detection, we used the built-in function (findpeaks) in MATLAB 2024b (MathWorks). To reduce dependency between detected peaks and increase their independence, we performed peak detection along lines separated by 6 wavelengths (∼0.6 mm) in the lateral directions. Specifically, for axial resolution measurements, the lines along which peaks were detected were spaced 6 wavelengths apart in both the X and Y axes. For lateral resolution measurements along the X (or Y) axis, the lines were spaced 6 wavelengths apart in the Y (or X) axis. Peaks were detected along an axis based on three criteria: 1) peak height must be greater than the average + 5 x standard deviation (std) of baseline signals for axial resolution, or average + 3 x std for lateral resolution; 2) peak prominence must be greater than the average + 5 x std for axial resolution, or average + 2 x std for lateral resolution; and 3) the minimum peak distance must be 0.15 mm for axial resolution, or 0.75 mm for lateral resolution. The baseline was extracted from regions without GV phantoms.

After computing the full-width-half-maximum (FWHM) of the detected peaks from all phantoms, the spatial resolution was calculated as the mean ± std of the FWHM values for each sequence (**Fig. 2k, Table 1**). The significance of differences in resolution among sequences was tested using the two-sided Mann-Whitney U test.

### Imaging GV-expressing brain tumor in mouse brains

GV-expressing U87 cells were prepared by genomic integration of second-generation acoustic reporter genes^10,13^. The implantation of these genetically engineered tumor cells and subsequent *in vivo* ultrasound imaging were conducted on three 5-month-old NSG male mice (Jackson Laboratory; denoted as mouse 1: **Fig. 3** and **Fig. 5b**, mouse 2: **Fig. S6**, and mouse 3: **Fig. S8a**) under protocols approved by the Institutional Animal Care and Use Committee of the California Institute of Technology.

1 × 10^5^ genetically engineered U87 cells were implanted at coordinates anterior-posterior (AP) –2 mm, medial-lateral (ML) +1.5 mm, and dorsal-ventral (DV) –3.5 mm. After allowing the tumor cells to grow for 6 days, we injected 150 μl of doxycycline through intraperitoneal (IP) injection for two consecutive days to induce the GV expression in the tumors. One day after the final doxycycline injection, the mice were anesthetized with 2-3% isoflurane, and a portion of the skull was removed to facilitate ultrasound imaging. After applying ultrasound gel, we initially imaged the mouse brain using a linear array probe (Verasonics, L22-14vX; center frequency 15.625 MHz) by taking xAM (angle = 19.5°, aperture size = 6.5 mm, PRF = 2 kHz)^31^ images of coronal slices with a 0.2 mm interval (except for mouse 3). The linear array probe was mounted on a stereotaxic instrument (Kopf Instruments), and its position was monitored using a digital display console. Next, the mouse brain was imaged with a matrix array probe using Takoyaki B-mode, AM and finally, BURST sequences. After collapsing the GVs, power Doppler images were acquired using the matrix array at the same location (except for mouse 2).

We used Napari^56^ for visualizing the volume images. The FOV of the doppler images was 9.6 × 9.6 mm, but the volume was cropped to match the FOV of Takoyaki AM and BURST. The dimensions of the volume image were (X, Y, Z) = 7.5 × 8.4 × 5.9 mm, with a depth range from 3.4 mm to 9.3 mm (**Fig. 3b**). For the anterior and lateral views, the volume images were half-cropped to exclude the skull side. The stacked 2D xAM (st-xAM) images were resized using bicubic interpolation from a voxel size of 50 × 200 × 1 *μ*m to 50 × 50 × 50 *μ*m so that it can be visualized in Napari with a reduced number of voxels. The volume size of the st-xAM image was 6.4 × 2.6 × 9 mm (**Fig. 3e**). We then manually registered the st-xAM image based on known probe locations relative to the bregma, patterns of GV signals and characteristic artifacts.

For CBR calculations, we drew a bounding box inscribed within the tumor to define the GV signal region for both Takoyaki AM and st-xAM images. The background region was defined as a bounding box of the same size, located at the contralateral counterpart of the tumor.

### Real-time monitoring of GV injections in the brain ventricles

The *in vivo* real-time monitoring experiment was performed on two 6-month-old C37bl/6 male mice (Jackson Laboratory, denoted as mouse 4: **Fig. 4** and **Fig. 5c**, and mouse 5: **Fig. S7** and **Fig. S8b**) under a protocol approved by the Institutional Animal Care and Use Committee of the California Institute of Technology. The mice were anesthetized with 2-3% isoflurane, and a portion of the skull was removed for imaging. A microliter syringe (Hamilton) was tilted at 45 degrees and targeted the right lateral ventricular of the mouse brain (AP –0.5 mm, ML +1 mm, DV –2 mm). A total of 4.5 *μ*L of GV solution (OD 14, equivalent to 1.6 nM^55^) was injected over 60 seconds at a rate of 75 nL/s using a micro syringe pump (World Precision Instrument), starting between the 11^th^ and 12^th^ frames. After applying ultrasound gel, a matrix probe was tilted at less than 10 degrees and placed without touching the micro-syringe.

Real-time imaging of the mouse brain was performed through the skull-removed window using the Takoyaki AM sequence, capturing data up to the 150^th^ frame for mouse 4 (300^th^ frame for mouse 5), which took approximately 10 mins (20 mins). Image reconstruction and necessary computations were performed using an Intel Xeon Gold 6136 CPU. The actual real-time visualization of live images was handled using Verasonics Vantage built-in functions, which displayed X, Y, and Z slices and volume images (not shown in this paper). Image reconstruction/visualization (∼2.8 sec/frame) and storage (∼1 sec/frame) were done after acquiring data for each frame, yielding a frame rate of 3.8 sec/frame. Power Doppler images were obtained before GV injection at the same location. After completing real-time monitoring, a BURST image was acquired using the same script as for real-time monitoring session, but with a high-pressure setting (Voltage ≥ 28 V).

For analysis, Volume Segmenter App in MATLAB 2024b (MathWorks) was used to manually label the lateral ventricles and the 3^rd^ ventricle. Regions affected by artifacts from the needle were excluded from the labels. Signal density was calculated by summing all signals within a labeled volume and dividing by the volume. 3D visualization of the stored images was done using Napari. The dimensions of the cropped Takoyaki AM image were (X, Y, Z) = 7.5 × 4.2 × 6.3 mm, with a depth range from 3.3 mm to 9.6 mm (**Fig. 4d**).

### Hydrophone measurements

Hydrophone measurements were performed in a water tank filled with degassed water, maintained using a water conditioner (Onda). The matrix probe was mounted on a motorized positioning stage (Velmex XSlide). A needle hydrophone (Onda HNR-0500) was submerged and fixed at the bottom of the tank, with its tip oriented perpendicularly toward the surface of the matrix probe. The hydrophone and Vantage 256 were connected to a digital oscilloscope (Keysight DSOX2004A) to measure acoustic pressure upon receiving trigger signals from the Vantage 256. Translating the probe across the field of interest, acoustic pressure was measured at each grid point. After averaging 256 received signals on the oscilloscope at each position, the averaged signal was sent to a connected computer, and the peak-to-peak pressure was calculated. For more accurate pressure measurements at the focus of the Takoyaki sequence (1^st^ TX aperture, **Fig. S4a**) within the Z = 7 mm plane, a fiber optic hydrophone system (Precision Acoustics) was used instead of the needle hydrophone. The pressure of plane waves was also measured at the same location using the fiber optic hydrophone (**Fig. S2b**).

## ACKNOWLEDGEMENTS

This research was supported by the National Institutes of Health (grants R01EB018975 and R01NS120828 to M.G.S.) and the Chan-Zuckerberg Initiative. M.G.S. is a Howard Hughes Medical Institute Investigator. Fig. 3a and Fig. 4a were created with Biorender.com.

## AUTHOR CONTRIBUTIONS

1. SL, CR, and MGS conceptualized the study. SL developed the Takoyaki sequence. CR developed the Doppler imaging sequence. DM prepared the GVs. Phantom preparation was carried out by SL and DW. SL conducted the *in vitro* experiments, and both CR and SL performed the *in vivo* experiments. Ultrasound data processing, analysis, and visualization were undertaken by SL. CR and MGS supervised the research. The original draft of the manuscript was written by SL, and SL, CR, DW, and MGS contributed to reviewing and editing the manuscript.

## DATA AND MATERIALS AVAILABAILITY

The ultrasound sequence scripts and data analysis codes used to generate key figures and results will be shared in a publicly accessible GitHub repository. Raw data are available from the corresponding authors upon reasonable request.

## Supplementary Materials

**Figure S1:**
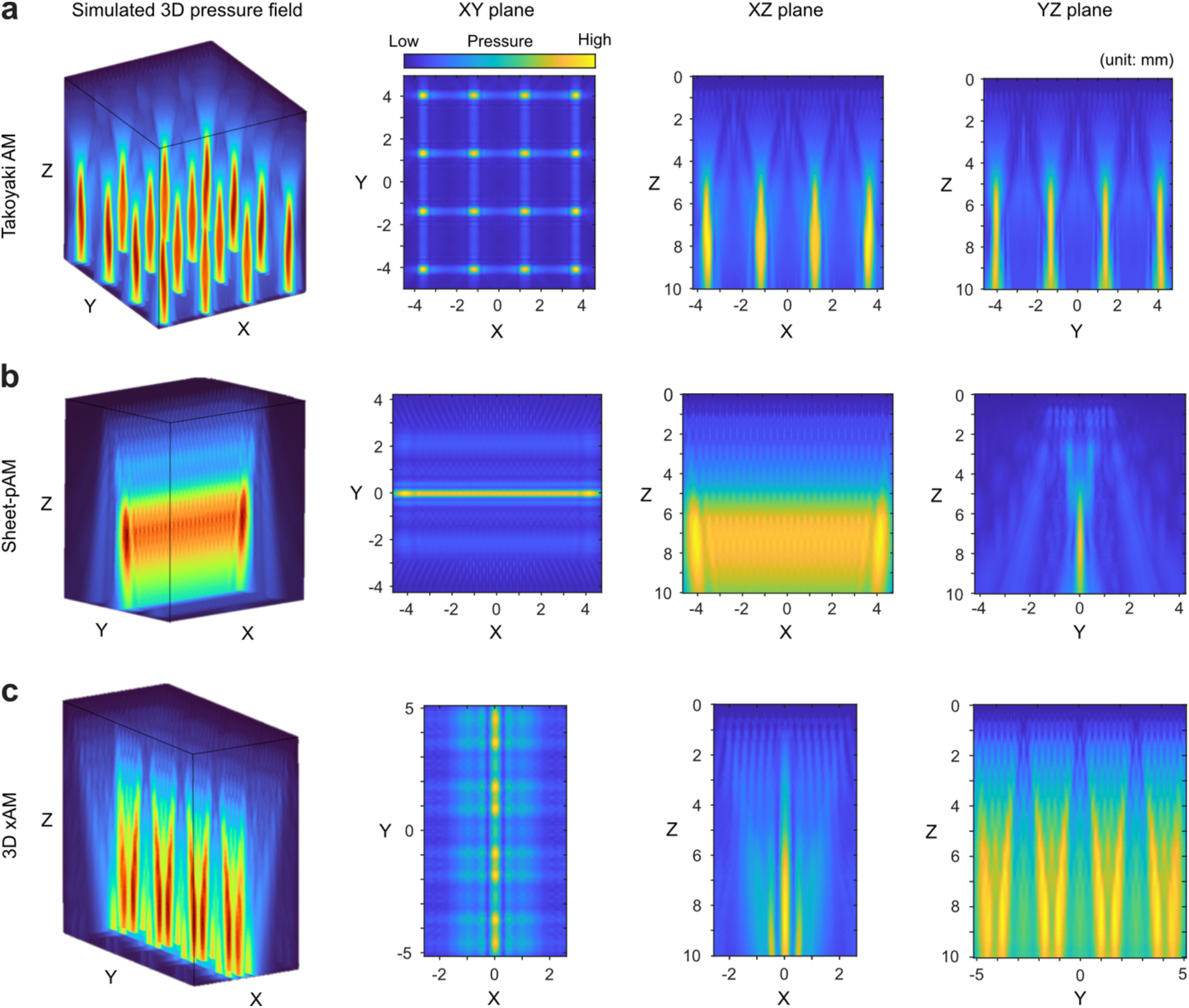
Simulated pressure fields. Pressure fields of Takoyaki (**a**), Sheet-pAM (**b**), and 3D xAM (**c**) simulated with TXPD in Verasonics Vantage software, based on the delays for full-amplitude shown in **Fig. 2a**. XY planes are extracted from Z = 7 mm. XZ planes of Takoyaki, Sheet-pAM, and 3D xAM pressure fields correspond to Y = 1.33 mm, Y = 0, and Y = 0.99 mm, respectively. YZ planes of Takoyaki, Sheet-pAM, and 3D xAM pressure fields correspond to X = 1.18 mm, X = 0, and X = 0, respectively.

**Figure S2:**
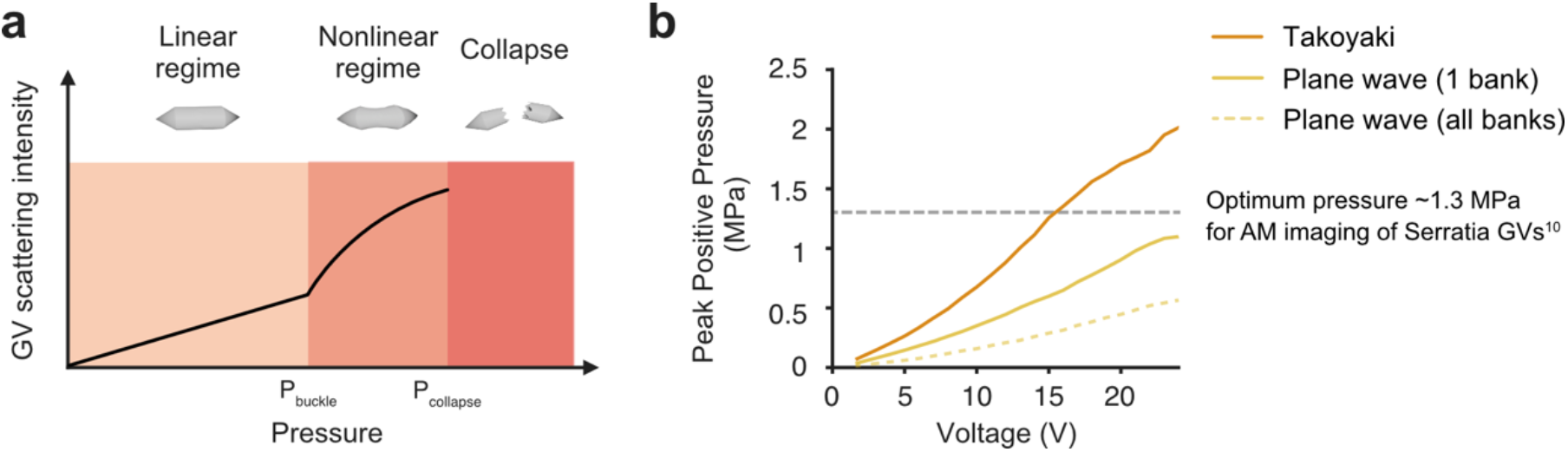
Pressure ranges of Takoyaki and plane-wave-based sequences. **a**. Illustration of different GV behaviors as a function of ultrasound pressure. Below the buckling threshold (P_buckle_), GVs exhibit linear scattering. Between P_buckle_ and the collapse threshold (P_collapse_), GVs begin to buckle, producing nonlinear responses suitable for AM imaging. Above P_collapse_, GVs collapse and generate strong, transient signals that can be isolated from the background using the BURST paradigm. **b**. Peak positive pressure of the Takoyaki sequence and plane waves emitted using one bank^29^ and all banks^57^, plotted against the input voltage of the MMA transducer. For certain types of GVs (e.g. Serratia GVs^10^), the pressure produced by plane waves is not enough to reach the optimum pressure for AM imaging. The pressures were measured at one of the focal points (Z = 7 mm) of the Takoyaki transmission shown in **Fig. S4a**, using a fiber optic hydrophone.

**Figure S3:**
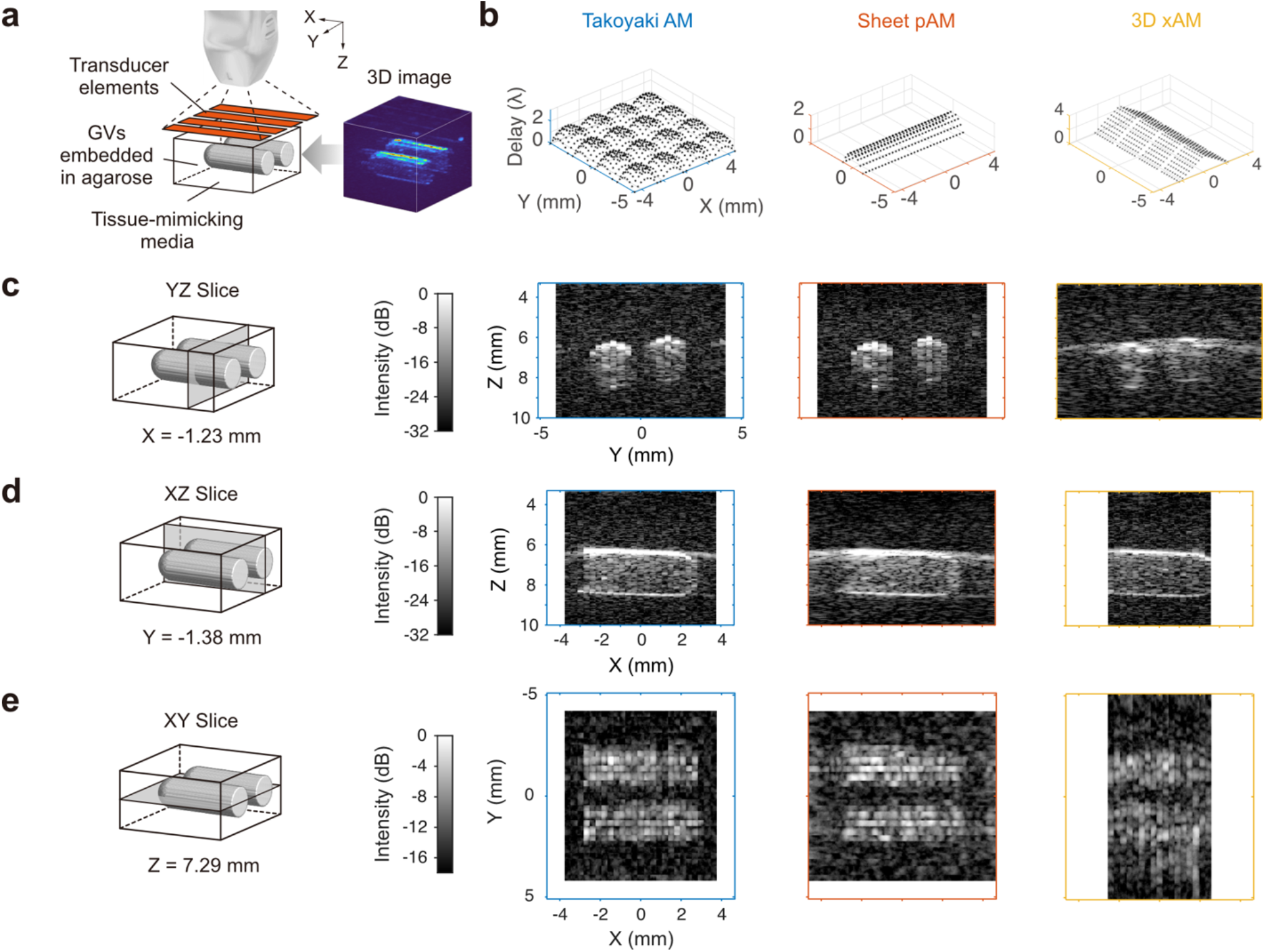
AM images of 90-degree rotated phantoms. **a**. Experiment setup for imaging tissue-mimicking GV phantoms. The orange parallelograms represent the four banks of transducer elements. The axis of the wells was aligned with the X axis, and the phantom location was adjusted so that its center was situated at Z = 7 mm. The 3D image of the phantom (OD = 6.5) acquired with Takoyaki AM is shown on the right. **b**. Representative delay laws of each sequence. **c-e**. Sliced phantom images acquired using different ultrasound sequences. For XY slices, five slices are averaged. The outer colored lines surrounding the sliced images indicate the maximum FOV. The color scale of the images is adjusted to match background levels across sequences.

**Figure S4:**
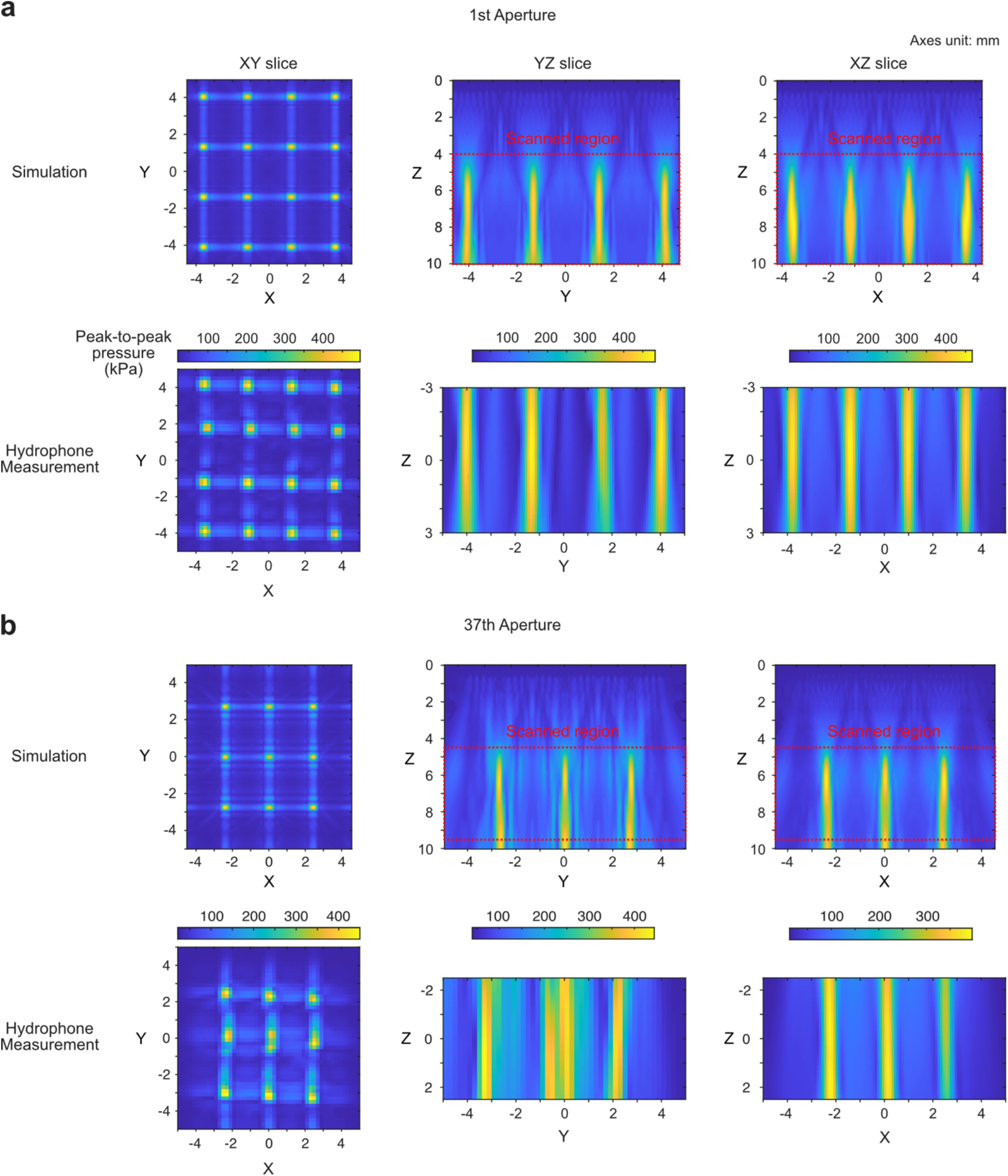
Measured pressured fields of Takoyaki sequence. Pressure fields of the Takoyaki sequence (focus = 7 mm) measured with a needle hydrophone. Their simulated pressure fields (ground truths) are shown in the top row of each panel. The coordinates of the hydrophone are centered at (0, 0, 7) mm in the simulation coordinate system. Red dotted boxes represent the regions scanned by the hydrophone. **a**. The first transmit aperture generates a (4 × 4) mesh of focused ultrasound. The XY, YZ, and XZ slices correspond to positions Z = 7 mm, X = 1.18 mm, and Y = 1.33 mm, respectively. While the focal points remain well-defined, deviations in their locations imply misalignment among the transducer element banks. This type of misalignment – where between-bank misalignment is more pronounced than with-bank misalignment – was also noted or assumed in previous element position calibration study for similar matrix array probe^42^. **b**. The 37^th^ transmit aperture emits a (3 × 3) mesh of focused ultrasound at the center of the FOV, with all focal points located within gap areas. The XY, YZ, and XZ slices correspond to positions Z = 7 mm, X = 0 mm, and Y = 0 mm, respectively. The pressure field at the Y = 0 mm plane is disrupted, likely due to the element bank misalignment. Combined with inherent gaps in the matrix array, which can hinder uniform pressure field application along the Y axis, these bank misalignments appear to cause the weakening of signals around Y = 0 mm.

**Figure S5:**
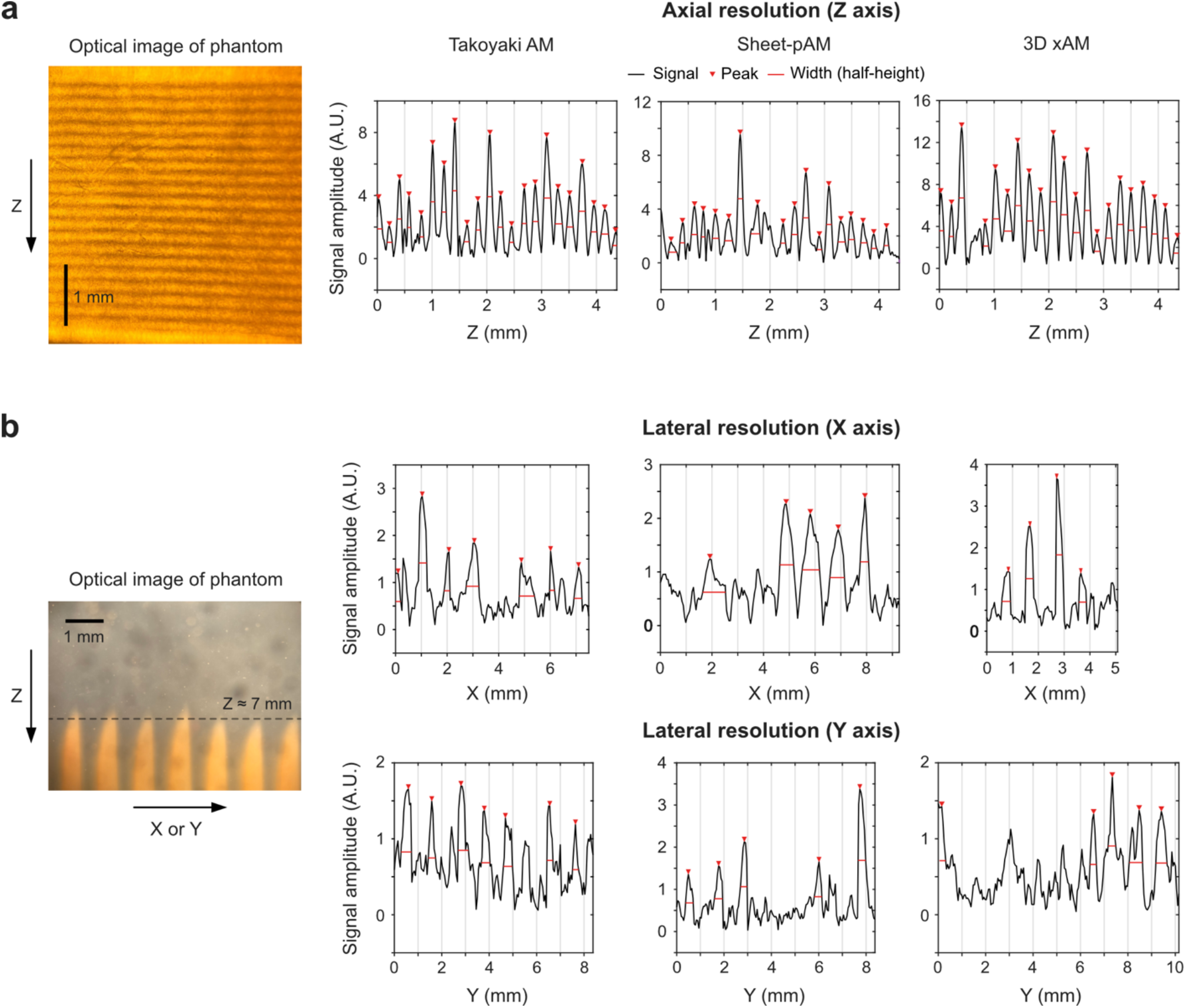
Spatial resolution measurements using phantoms with GV stripes. Detected peaks and their full-width-half-maximum (FWHM) are marked with red triangles and red lines, respectively. **a**. Axial (Z axis) resolution measurements. Phantoms have GV stripes with an interval of 0.2 mm. For each sequence, an example of signals along the Z axis is shown. **b**. Lateral (X and Y axes) resolution measurements. Phantoms have a saw-tooth pattern of GVs with the interval of 1 mm, and we chose the plane of Z = ∼7 mm (sharp regions of the saw-tooth) for FWHM analysis. The top and bottom rows correspond to the X and Y directions, respectively.

**Figure S6:**
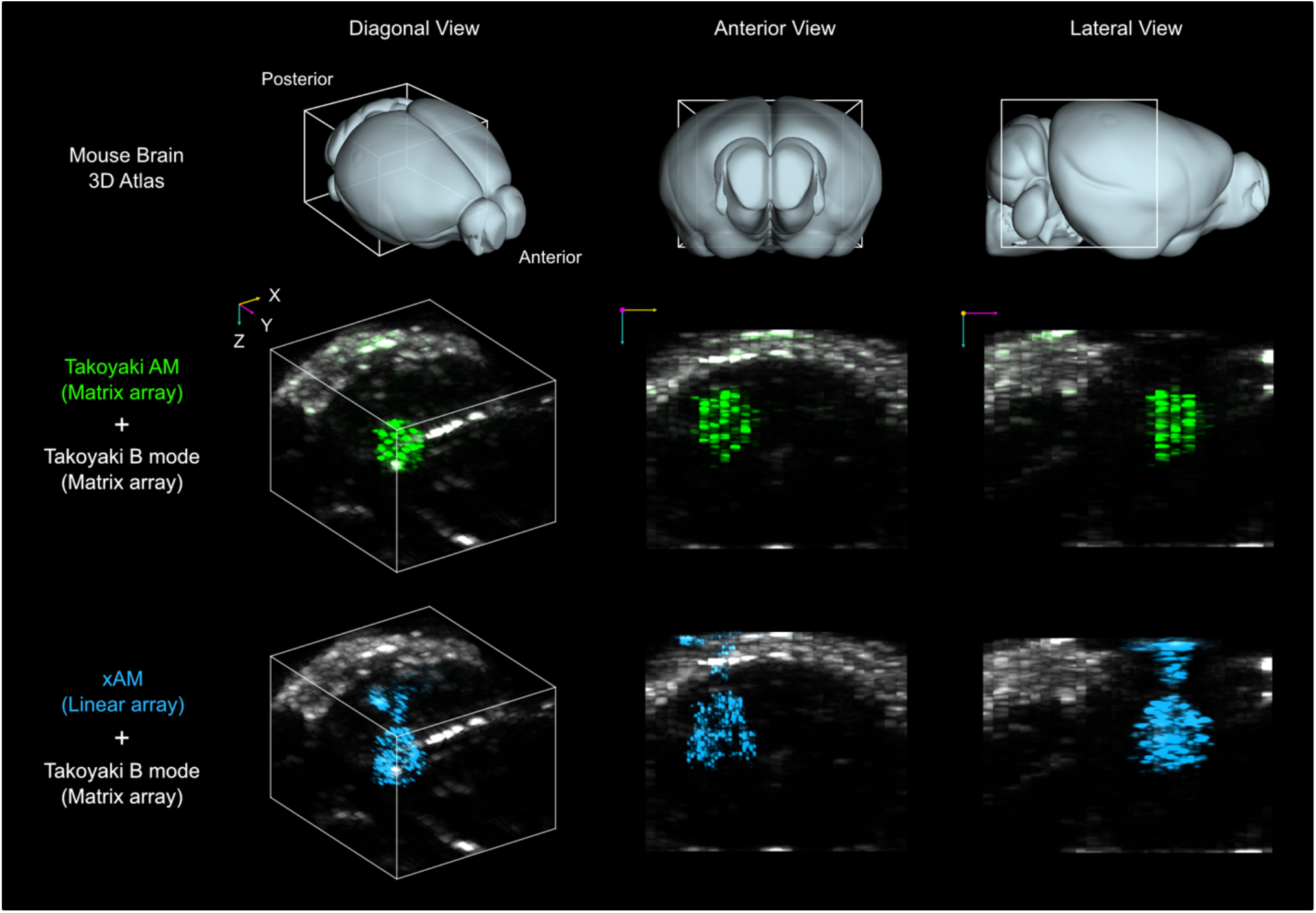
Additional example of imaging genetically labeled tumor in mouse brain with the Takoyaki AM and 2D xAM. Both images are overlaid with Takoyaki B-mode. The size of the left column’s volume images is (X, Y, Z) = 7.5 × 8.4 × 6.3 mm, with a depth range from 3.4 mm to 9.7 mm. The white box on the brain atlas indicates the imaging regions. Colored arrows represent the axes, with lengths of 1 mm along their corresponding directions. CBRs for Takoyaki AM and st-xAM were 11.27 and 10.33 dB, respectively.

**Figure S7:**
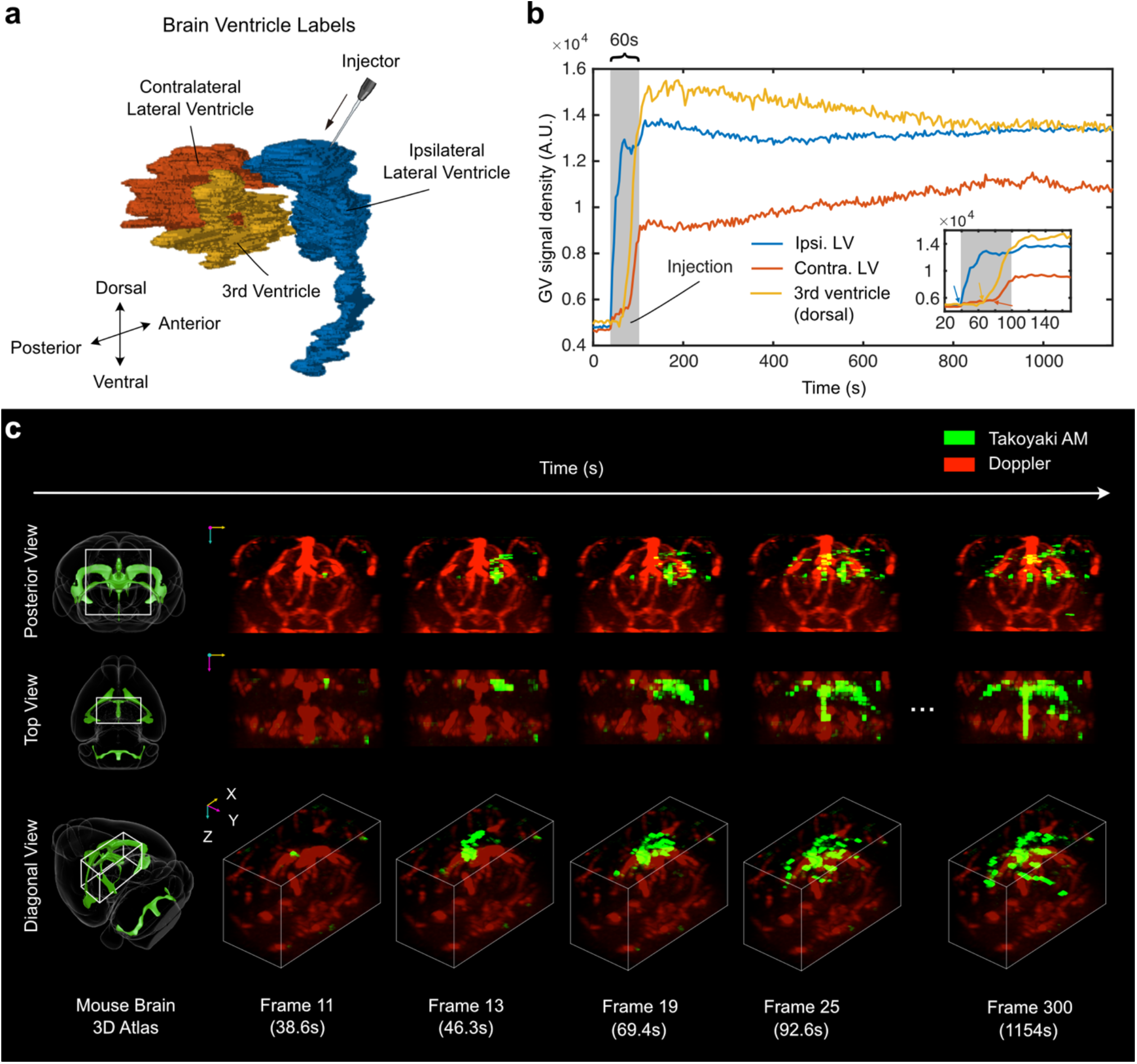
Additional example of monitoring GV propagation in brain ventricles. **a**. Manually drawn outlines of brain ventricles. **b**. Temporal evolution of GV signal density. GV injection (gray shaded area) begins between the frame 11 (38.6s) and 12 (42.5s) **c**. Selected frames from stored real-time images, visualized with Napari. Takoyaki AM images (green) are overlaid on power Doppler images (red). Volume images are half-cropped to exclude the skull side. The dimensions of the cropped Takoyaki AM image are (X, Y, Z) = 7.5 × 4.2 × 6.0 mm, with a depth range from 3.3 mm to 9.3 mm. The sampling interval was approximately 3.8 sec. The 3D Allen Mouse Brain Atlas in the leftmost column illustrates brain orientations. White bounding boxes show the imaging location. Colored arrows have a length of 1 mm along their corresponding directions.

**Figure S8:**
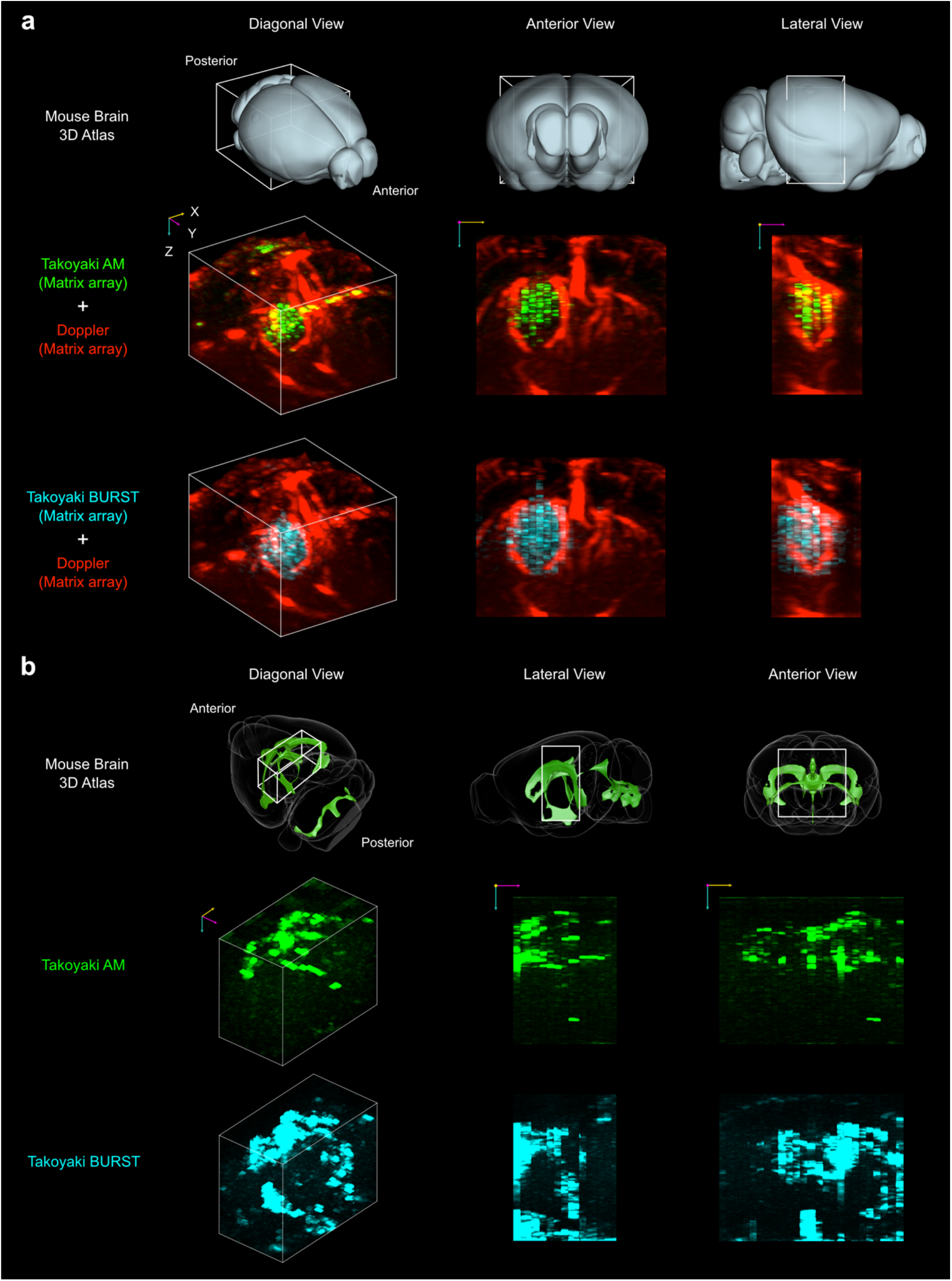
Additional examples of Takoyaki AM and BURST images of (a) genetically labeled tumors and (b) brain ventricles. **a**. The dimensions of the volume images (left column) is (X, Y, Z) = 7.5 × 8.4 × 5.9 mm, with a depth range from 3.4 mm to 9.3 mm. The skull areas are excluded in the anterior and lateral views. **b**. Takoyaki AM images from the last frame of real-time recording session (top row) and Takoyaki BURST images (bottom row) acquired after real-time recording. The dimensions of the volume images (left column) are (X, Y, Z) = 7.5 × 4.2 × 6 mm, with a depth range from 3.3 mm to 9.3 mm. The skull areas are excluded in all views. The white box indicates the approximate imaging regions. Colored arrows represent the axes, with lengths of 1 mm.

## Description of supplementary videos

**Video 1**: 3D images of GV phantoms acquired with Takoyaki AM, Sheet-pAM, and 3D xAM, corresponding to **Fig. 2c-e**. Magenta, yellow and cyan arrows represent the X, Y, and Z axes, respectively.

**Video 2**: 3D images of genetically labeled tumors in the mouse brain, corresponding to **Fig. 3d-e**. Takoyaki AM and st-xAM images are overlaid with a power Doppler image. Magenta, yellow and cyan arrows represent the X, Y, and Z axes, respectively.

**Video 3**: Real-time recording of nanoparticle transport within mouse brain ventricles, corresponding to **Fig. 4d**. The frame rate is accelerated 20 times. Yellow, magenta and cyan arrows represent the X, Y, and Z axes, respectively. The length of each arrow corresponds to 1 mm.

**Video 4**: 3D images of genetically labeled tumors and brain ventricles acquired with Takoyaki AM and BURST, corresponding to **Fig. 5b-c**. Magenta, yellow and cyan arrows represent the X, Y, and Z axes, respectively.

## REFERENCES

1. Huang, Q. & Zeng, Z. A Review on Real-Time 3D Ultrasound Imaging Technology. BioMed Res. Int. 2017, 6027029 (2017).

2. Yusefi, H. & Helfield, B. Ultrasound Contrast Imaging: Fundamentals and Emerging Technology. Front. Phys. 10, (2022).

3. Frinking, P., Segers, T., Luan, Y. & Tranquart, F. Three Decades of Ultrasound Contrast Agents: A Review of the Past, Present and Future Improvements. Ultrasound Med. Biol. 46, 892–908 (2020).

4. Leon, A. de et al. Contrast enhanced ultrasound imaging by nature-inspired ultrastable echogenic nanobubbles. Nanoscale 11, 15647–15658 (2019).

5. Sheeran, P. S. & Dayton, P. A. Phase-Change Contrast Agents for Imaging and Therapy. Curr. Pharm. Des. 18, 2152– 2165 (2012).

6. Shapiro, M. G. et al. Biogenic gas nanostructures as ultrasonic molecular reporters. Nat. Nanotechnol. 9, 311–316 (2014).

7. Walsby, A. E. Gas vesicles. Microbiol. Rev. 58, 94–144 (1994).

8. Bourdeau, R. W. et al. Acoustic reporter genes for noninvasive imaging of microorganisms in mammalian hosts. Nature 553, 86–90 (2018).

9. Farhadi, A., Ho, G. H., Sawyer, D. P., Bourdeau, R. W. & Shapiro, M. G. Ultrasound imaging of gene expression in mammalian cells. Science 365, 1469–1475 (2019).

10. Hurt, R. C. et al. Genomically mined acoustic reporter genes for real-time in vivo monitoring of tumors and tumor-homing bacteria. Nat. Biotechnol. 41, 919–931 (2023).

11. Buss, M. T., Zhu, L., Kwon, J. H., Tabor, J. J. & Shapiro, M. G. Probiotic acoustic biosensors for noninvasive imaging of gut inflammation. 2024.09.23.614598 Preprint at 10.1101/2024.09.23.614598 (2024).

12. Nyström, N. N. et al. Multiplexed Ultrasound Imaging of Gene Expression. 2024.10.30.621148 Preprint at 10.1101/2024.10.30.621148 (2024).

13. Shivaei, S. et al. Non-invasive imaging of cell-based therapies using acoustic reporter genes. 2024.11.01.621111 Preprint at 10.1101/2024.11.01.621111 (2024).

14. Jin, Z. et al. Ultrasonic reporters of calcium for deep tissue imaging of cellular signals. 2023.11.09.566364 Preprint at 10.1101/2023.11.09.566364 (2023).

15. Lakshmanan, A. et al. Acoustic biosensors for ultrasound imaging of enzyme activity. Nat. Chem. Biol. 16, 988–996 (2020).

16. Le Floc’h, J. et al. In vivo Biodistribution of Radiolabeled Acoustic Protein Nanostructures. Mol. Imaging Biol. 20, 230–239 (2018).

17. Wang, G. et al. Surface-modified GVs as nanosized contrast agents for molecular ultrasound imaging of tumor. Biomaterials 236, 119803 (2020).

18. Ling, B. et al. Biomolecular Ultrasound Imaging of Phagolysosomal Function. ACS Nano 14, 12210–12221 (2020).

19. Wang, Y. et al. Modification of PEG reduces the immunogenicity of biosynthetic gas vesicles. Front. Bioeng. Biotechnol. 11, (2023).

20. Ling, B. et al. Gas Vesicle–Blood Interactions Enhance Ultrasound Imaging Contrast. Nano Lett. 23, 10748–10757 (2023).

21. Ling, B. et al. Truly Tiny Acoustic Biomolecules for Ultrasound Imaging and Therapy. Adv. Mater. 36, 2307106 (2024).

22. Provost, J. et al. 3D ultrafast ultrasound imaging in vivo. Phys. Med. Biol. 59, L1 (2014).

23. Provost, J. et al. 3-D ultrafast doppler imaging applied to the noninvasive mapping of blood vessels in Vivo. IEEE Trans. Ultrason. Ferroelectr. Freq. Control 62, 1467–1472 (2015).

24. Roux, E. et al. Experimental 3-D Ultrasound Imaging with 2-D Sparse Arrays using Focused and Diverging Waves. Sci. Rep. 8, 9108 (2018).

25. Heiles, B. et al. Ultrafast 3D Ultrasound Localization Microscopy Using a 32 \times 32 Matrix Array. IEEE Trans. Med. Imaging 38, 2005–2015 (2019).

26. Rabut, C. et al. 4D functional ultrasound imaging of whole-brain activity in rodents. Nat. Methods 16, 994 (2019). manuscript was written by SL, and SL, CR, DW, and MGS contributed to reviewing and editing the manuscript.

27. Sauvage, J. et al. 4D Functional Imaging of the Rat Brain Using a Large Aperture Row-Column Array. IEEE Trans. Med. Imaging 39, 1884–1893 (2020).

28. Yu, J., Yoon, H., Khalifa, Y. M. & Emelianov, S. Y. Design of a Volumetric Imaging Sequence Using a Vantage-Ultrasound Research Platform Multiplexed with a 1024-Element Fully-Sampled Matrix Array. IEEE Trans. Ultrason. Ferroelectr. Freq. Control 67, 248–257 (2020).

29. Chavignon, A. et al. 3D Transcranial Ultrasound Localization Microscopy in the Rat Brain With a Multiplexed Matrix Probe. IEEE Trans. Biomed. Eng. 69, 2132–2142 (2022).

30. Maresca, D. et al. Nonlinear ultrasound imaging of nanoscale acoustic biomolecules. Appl. Phys. Lett. 110, 073704 (2017).

31. Maresca, D., Sawyer, D. P., Renaud, G., Lee-Gosselin, A. & Shapiro, M. G. Nonlinear X-Wave Ultrasound Imaging of Acoustic Biomolecules. Phys. Rev. X 8, 041002 (2018).

32. Rabut, C. et al. Ultrafast amplitude modulation for molecular and hemodynamic ultrasound imaging. Appl. Phys. Lett. 118, 244102 (2021).

33. Sawyer, D. P. et al. Ultrasensitive Ultrasound Imaging of Gene Expression with Signal Unmixing. Nat. Methods 18, 945–952 (2021).

34. Cherin, E. et al. Acoustic Behavior of Halobacterium salinarum Gas Vesicles in the High-Frequency Range: Experiments and Modeling. Ultrasound Med. Biol. 43, 1016–1030 (2017).

35. Zhang, S. et al. The Vibration Behavior of Sub-Micrometer Gas Vesicles in Response to Acoustic Excitation Determined via Laser Doppler Vibrometry. Adv. Funct. Mater. 30, 2000239 (2020).

36. Rasmussen, M. F., Christiansen, T. L., Thomsen, E. V. & Jensen, J. A. 3-D imaging using row-column-addressed arrays with integrated apodization – part i: apodization design and line element beamforming. IEEE Trans. Ultrason. Ferroelectr. Freq. Control 62, 947–958 (2015).

37. Christiansen, T. L. et al. 3-D imaging using row–column-addressed arrays with integrated apodization— part ii: transducer fabrication and experimental results. IEEE Trans. Ultrason. Ferroelectr. Freq. Control 62, 959–971 (2015).

38. Bouzari, H. et al. Imaging Performance for Two Row–Column Arrays. IEEE Trans. Ultrason. Ferroelectr. Freq. Control 66, 1209–1221 (2019).

39. Heiles, B. et al. Nonlinear sound-sheet microscopy: imaging opaque organs at the capillary and cellular scale. 2024.07.31.605825 Preprint at 10.1101/2024.07.31.605825 (2024).

40. Bernal, M., Cunitz, B., Rohrbach, D. & Daigle, R. High-frame-rate volume imaging using sparse-random-aperture compounding. Phys. Med. Biol. 65, 175002 (2020).

41. Mallart, R. & Fink, M. Improved imaging rate through simultaneous transmission of several ultrasound beams. In New Developments in Ultrasonic Transducers and Transducer Systems vol. 1733 120–130 (SPIE, 1992).

42. McCall, J. R., Chavignon, A., Couture, O., Dayton, P. A. & Pinton, G. F. Element Position Calibration for Matrix Array Transducers with Multiple Disjoint Piezoelectric Panels. Ultrason. Imaging 46, 139–150 (2024).

43. Wang, Z., Simoncelli, E. P. & Bovik, A. C. Multiscale structural similarity for image quality assessment. in The Thrity-Seventh Asilomar Conference on Signals, Systems & Computers, 2003 vol. 2 1398-1402 Vol.2 (2003).

44. Wu, D. et al. Biomolecular actuators for genetically selective acoustic manipulation of cells. Sci. Adv. 9, eadd9186 (2023).

45. Rabut et al., C. in preparation.

46. Wang, Q. et al. The Allen Mouse Brain Common Coordinate Framework: A 3D Reference Atlas. Cell 181, 936-953.e20 (2020).

47. Xie, L. et al. Sleep Drives Metabolite Clearance from the Adult Brain. Science 342, 373–377 (2013).

48. Kelley, D. H. Brain cerebrospinal fluid flow. Phys. Rev. Fluids 6, 070501 (2021).

49. Kelley, D. H. & Thomas, J. H. Cerebrospinal Fluid Flow. Annu. Rev. Fluid Mech. 55, 237–264 (2023).

50. Hughes, M. P. et al. AAV9 intracerebroventricular gene therapy improves lifespan, locomotor function and pathology in a mouse model of Niemann–Pick type C1 disease. Hum. Mol. Genet. 27, 3079–3098 (2018).

51. Taghian, T. et al. A Safe and Reliable Technique for CNS Delivery of AAV Vectors in the Cisterna Magna. Mol. Ther. 28, 411–421 (2020).

52. Hurt, R. C. et al. Plasmid from Articles – Genomically mined acoustic reporter genes for real-time in vivo monitoring of tumors and tumor-homing bacteria. Addgene https://www.addgene.org/browse/article/28229434/.

53. Demené, C. et al. Spatiotemporal Clutter Filtering of Ultrafast Ultrasound Data Highly Increases Doppler and fUltrasound Sensitivity. IEEE Trans. Med. Imaging 34, 2271–2285 (2015).

54. Eckart, C. & Young, G. The approximation of one matrix by another of lower rank. Psychometrika 1, 211–218 (1936).

55. Lakshmanan, A. et al. Preparation of biogenic gas vesicle nanostructures for use as contrast agents for ultrasound and MRI. Nat. Protoc. 12, 2050–2080 (2017).

56. Ahlers, J. et al. napari: a multi-dimensional image viewer for Python. Zenodo 10.5281/zenodo.8115575 (2023).

57. Xing, P. et al. Towards Transcranial 3D Ultrasound Localization Microscopy of the Nonhuman Primate Brain. Preprint at 10.48550/arXiv.2404.03547 (2024).

